# MYC and RNA Polymerase II Binding Near Transcriptional End Sites Regulate the Expression of Functionally-Related Genes

**DOI:** 10.64898/2026.06.22.733817

**Authors:** Colin M. Henchy, Huabo Wang, Edward V. Prochownik

## Abstract

MYC oncoprotein binding at promoters and enhancers influences RNA polymerase II (RNAPII)-driven gene expression. Numerous genes also bind MYC near their transcriptional end sites (TESs). This often allows direct promoter-TES contact via looping and further regulates total and “read-through” transcription that extends beyond standard termination sites. We aimed here to better clarify the rules governing TES-associated MYC and/or RNAPII binding cross-talk in human and murine cells. Using ChIPseq and RNAseq datasets from the ENCODE portal and elsewhere, MYC and RNAPII binding profiles were found to differ around TESs and transcriptional start sites (TSSs). Variations in E box flanking sequences likely accounted for the somewhat lower affinities of MYC for TES-associated sites. Motifs for numerous other transcription factors were also observed to cluster non-randomly and in close proximity to MYC and RNAPII binding site peak summits. On average, genes with TES-proximal MYC or RNAPII sites were more highly expressed than those without, although co-binding tended to be suppressive. Both normal and neoplastic proliferative stimuli altered the MYC and RNAPII binding patterns of many genes, indicating that “category switching” was common, subject to disparate external signals and often reversible. Functionally related gene sets with high levels of read-through transcription were uniformly marked by significant amounts of TES-associated MYC and/or RNAPII binding. These findings indicate that, both independently and together, MYC and RNAPII binding near TESs dynamically impact total and read-through transcription while also coordinating the expression of many common purpose gene sets.

**Significance Statement:** Binding of the MYC oncoprotein around transcriptional end sites (TESs) of genes can influence both total and “readthrough” transcription that extends beyond normal termination points. We have now studied how RNA polymerase II (RNAPII) cross-talks with MYC to regulate these processes and their consequences. We found that binding of these factors around TESs cooperates with their binding at transcriptional start sites (TSSs) and increases both absolute and read-through transcription. MYC and/or RNAPII binding around TSSs and TESs is often transient and reversible in response to both normal and neoplastic growth-promoting stimuli. TES-associated MYC and/or RNAPII binding thus not only influence the quantity and quality of gene expression but also mark gene subsets that participate in common pathways and functions.

## Introduction

MYC is a classic bHLH-Zip-type transcription factor (TF) that is evolutionarily conserved across all metazoan species, is highly expressed by proliferating normal cells, and is deregulated in most cancers (1–3). Its role in transcriptional activation requires mandatory heterodimerization with another bHLH-Zip protein, MAX, in order to bind “E box” DNA elements that usually reside in the proximal promoters of its target genes (2, 4, 5). This binding then initiates a complex and coordinated series of events that include local chromatin acetylation and relaxation, MYC’s interaction with RNA polymerase II (RNAPII) or other RNAPs and the recruitment of various co-factors that initiate *de novo* transcription and relieve transcriptional pausing (2, 6–9). Although it has been proposed that MYC amplifies transcription on a global scale, the most highly de-regulated gene targets tend to be those involved in specific growth-related functions such as cell cycle control, metabolism, ribosomal biogenesis, translation and mRNA splicing (2, 4, 6, 10–12).

MYC target genes can also be negatively regulated when MYC-MAX heterodimers are displaced from E boxes by competing heterodimers comprised of MAX and members of the bHLH-Zip “MXD family” (MXD1-4, MNT, and MGA) (2, 4, 13). The ensuing de-acetylation and re-compaction of promoter-associated chromatin facilitate a return to the genes’ original and less transcriptionally permissive states (2, 4, 13). The competition between MYC and MXD members for MAX is particularly prominent during periods of cellular quiescence or differentiation when MYC levels are low and/or declining and MXD member levels are high and/or increasing, often in tissue-specific ways. Other forms of MYC target gene transcriptional suppression are indirect and involve inhibitory interactions between MYC-MAX heterodimers and the positively-acting TFs SP1/3 and MIZ1/ZBTB17 (hereafter ZBTB17), which bind their own unique and eponymous consensus motifs (2, 14–16). The number of MYC-responsive target genes directly correlates with MYC levels and inversely with MXD member levels (2, 4, 11, 17). This has led to the realization that some MYC-regulated genes with low-affinity binding sites are “pathologic” targets whose expression is altered primarily or exclusively in response to the aberrantly high levels of MYC present in some cancers and whose contributions to oncogenesis, while essential, remain incompletely understood (18–21).

MYC-MAX heterodimers can also bind enhancers, particularly when MYC is over-expressed by tumors. Among the consequences of this are a more open chromatin environment within the enhancer, the transcription of “eRNAs” originating from this region and direct and cooperative contacts with promoter-localized MYC-MAX that further elevate and customize target gene transcription (22–26).

We have recently described and characterized a previously unappreciated form of gene regulation mediated by MYC and/or MAX binding at the 3’-ends of ∼15-20% of all protein-coding and long non-coding (lnc) genes (27). Unlike promoter-associated MYC, which tends to cluster in close proximity to transcriptional start sites (TSSs), or enhancer-associated MYC, which binds at much greater and less predictable distances from TSSs, MYC’s localization at the 3’-ends of genes broadly centers around transcriptional end sites (TESs) (27). These regions are also highly enriched for RNAPII and numerous other TFs whose collective binding patterns readily distinguish them from TSSs and enhancers (27). Like promoters, MYC-bound TES-proximal regions display transcriptionally conducive acetylation and methylation of neighboring histones and high nuclease susceptibility. In many cases, they also directly contact MYC-bound TSSs and/or enhancers via looping. MYC binding at TESs also marks genes whose functions differ from those lacking such binding (27). Finally, in addition to regulating basal gene expression, TES-associated MYC binding can also regulate “read-through” or “downstream of gene” (DoG) transcription that tends to be stress-induced and exerts control over nuclear-cytoplasmic transport, translation and the transcriptional control of adjacent downstream genes (28–32).

The identification of TES-associated MYC binding in close proximity to numerous other TFs and TF binding motifs and the participation of such binding in both quantitative and qualitative aspects of gene regulation suggest that these functions are governed by an as yet poorly understood set of rules. Among the conditions that might be expected to define these are the identities of the MYC-associated factors, whether they (and MYC itself) also bind to TSSs and the interactions between these sites (7, 33–35). To begin to understand what is likely a complex, interactive and dynamic regulatory process involving both common and tissue-specific factors, we have compared the properties of genes associated with MYC and/or RNAPII binding at TSSs and/or TESs. Our findings reveal that this simple type of categorization is dynamic and predictive of gene expression levels, their functions and the degree to which they engage in DoG transcription.

## Results

### TSSs and TESs display distinct profiles of MYC and RNAPII DNA binding

Both direct (E box-associated) and indirect (SP1/3- and ZBTB17-associated) site-specific DNA binding by MYC are MAX-dependent, occur at promoters and enhancers, and often involve RNAPII (2, 7, 11, 14–16, 36–41). However, the relationships between MYC and RNAPII at TES-proximal sites have not been previously investigated. Using five human and two murine cell lines for which MYC and RNAPII ChIPseq profiles and RNAseq results were publicly available from the ENCODE database (https://www.encodeproject.org/), we compiled an unbiased, genome-wide list of MYC and RNAPII binding sites residing in proximity to TESs and compared these profiles to those at TSSs. Five of the cell lines (HepG2, K562, A549, MEL and CH12.lx) were cancer-derived and two (H1 and HUVEC) originated from non-transformed human embryonic stem cells and umbilical vein endothelial cells, respectively. Basal MYC levels among these seven cell lines varied over a nearly six-fold range, with murine erythroleukemia (MEL) cells expressing the highest levels and H1 cells expressing the lowest (Supplementary Table 1). Because no significant differences were observed in the binding patterns among protein-coding and lnc genes, they were combined to maximize statistical power (27, 42). As done previously, we excluded genes <2.5 kb in length, which prevented the unequivocal assignment of MYC and RNAPII ChIP peaks to TSSs or TESs. Also excluded were genes arranged sufficiently close to one another and in head-to-tail configuration that led to ambiguous binding site assignments (27). MYC and RNAPII binding peak summits located within 2.5 kb of two TSSs or two TESs of adjacent genes arranged in head-to-head or tail-to-tail configuration, respectively, were assigned to both genes as both TFs have been shown to affect the expression of each member of such closely-spaced pairs (27, 43–45).

We first compared the total MYC and RNAPII binding profiles for genes that associated with these factors around TSSs and/or TESs. As we previously reported, 61.2% +/- 2.5% of all 5’ MYC ChIPseq footprints mapped to within +/- 500 bp of TSSs with binding downstream of these sites (54.1% +/- 1.1%) being slightly preferred to binding upstream (45.9% +/- 1.1%, p=0.03) (Figure 1A, Supplementary Table 2). This agreed with previous findings regarding the distribution of these sites as well as those for Sp1 and ZBTB17 (5, 27, 46, 47). RNAPII ChIPseq footprints overlapped significantly with those for MYC, with an average of 67.6% +/-2.5% also residing within 500 bps of TSSs (Supplementary Table 2). Relative to MYC footprints, more RNAPII footprints resided downstream of TSSs (62.8% +/- 0.9% vs. 54.1% +/- 1.1%, p=0.015) (Supplementary Table 3). This was consistent with the often bi-modal distribution of RNAPII, with the more downstream of the peaks centering ∼30-100 bp from TSSs and coinciding with sites of transcriptional pausing (Supplementary Table 4) (48, 49). On average, MYC occupied more than four times as many sites in transformed cell lines as in non-transformed cell lines (Figure 1A). This may reflect the 1.8-fold higher mean levels of MYC in the former group (Supplementary Table 1) and its occupation of more low-affinity sites as well as the more transcriptionally accessible landscape of cancer cells in general (18–23, 27, 50).

**Figure 1.**
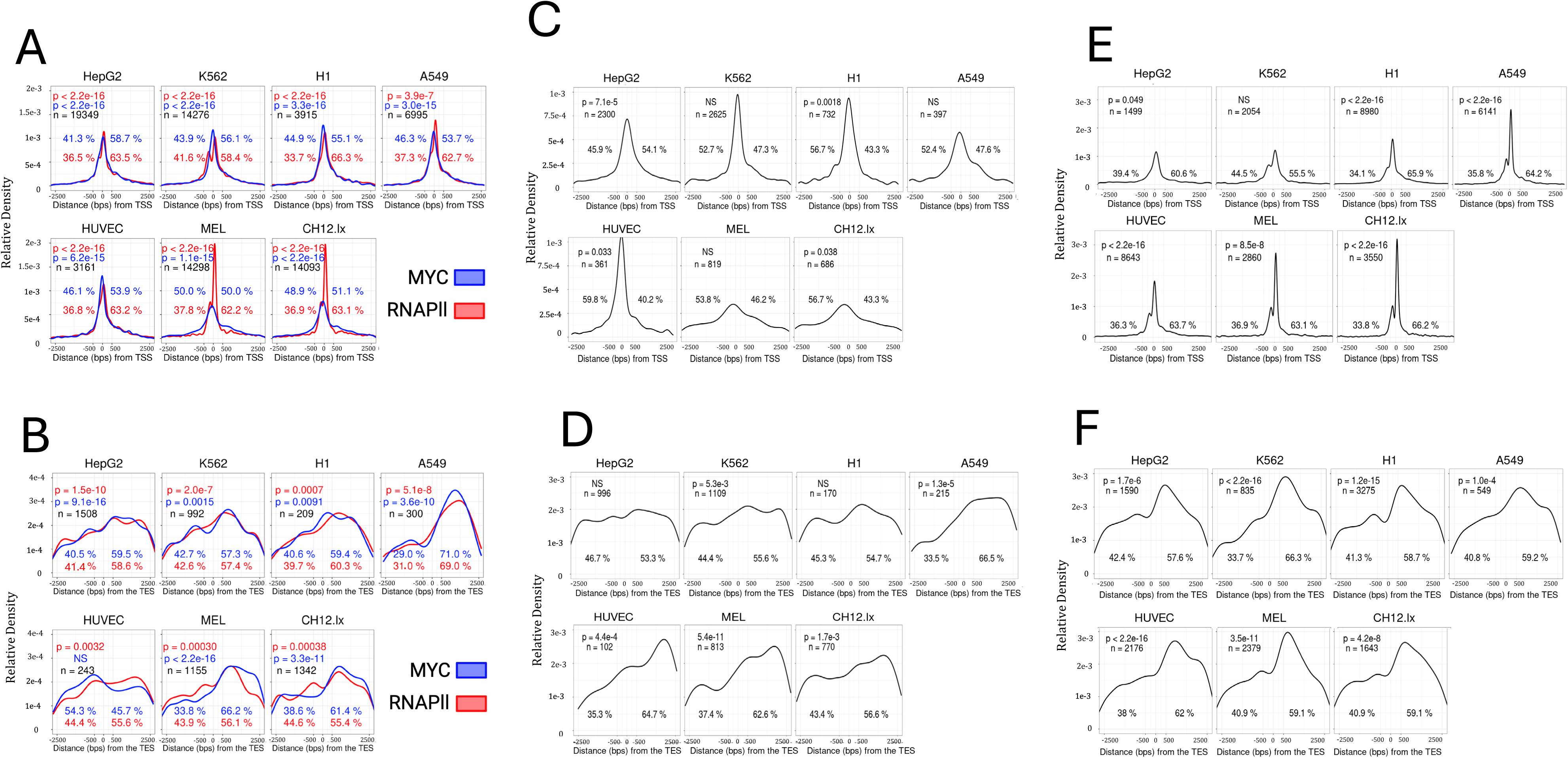
Binding of MYC and RNAPII at TSSs and TESs in seven human and murine cell lines. (A). Distributions of MYC and RNAPII binding sites around TSSs of genes with at least one site for either factor. In this panel and all others, p values, determined by the Wilcoxon Signed Rank Test, indicate the likelihood that the factor would otherwise randomly center precisely at the TSS (arbitrarily defined as “0”). Numbers in black beneath these p values indicate the total number of binding sites that were included in each analysis. Because each ChIPSeq peak was included, some genes were counted more than once if they had multiple binding sites (27). Red and blue numbers adjacent to the profiles indicate the percentage of all peaks that mapped upstream or downstream of the TSS. (B). Binding patterns of MYC and RNAPII around TESs of genes with at least one site for either factor. As in A, some genes were included more than once to permit multiple binding site assignments. (C). MYC binding patterns at TSSs of genes that bound only MYC. As in A and B, some genes were included more than once if they contained multiple binding sites. (D). MYC binding patterns at TESs of genes that bound only MYC. As described above, some genes were included more than once if they contained multiple binding sites. (E). RNAPII binding patterns at TSSs of genes that bound only RNAPII. As described above, some genes were included more than once if they contained multiple binding sites (F). RNAPII binding patterns at TESs of genes that bound only RNAPII. As described above, some genes were included more than once if they contained multiple binding sites

In contrast to the tight clustering of MYC and RNAPII binding sites around TSSs, the distribution of these factors around TESs, while continuing to be associated with significant overlap, was more diffuse (Figure 1B). On average, only 39.0% +/- 0.6% of MYC peak summits localized to within +/- 1kb of TESs as did only 41.7% +/- 0.6% of those for RNAPII (Supplementary Table 2). The fraction of MYC binding downstream of TESs was not significantly greater than at TSSs (60.0% +/- 3.0% vs. 54.1% +/- 1.1%, p=0.11) nor was the fraction of RNAPII binding downstream of TESs (58.9% +/- 1.8% vs. 62.8% +/- 0.9% p=0.16) (Supplementary Table 3). More striking, however, was that TES-associated footprint peak summits were, on average, located further downstream than were TSS footprint peak summits, were more variable, and in some cases were distributed bi- or even tri-modally (Figure 1 and Supplementary Table 5).

From the above groups, we more closely analyzed MYC ChIPseq footprints at TSS- or TES-proximal sites in the absence of demonstrable RNAPII co-binding (Figure 1C and D, respectively). On average, these profiles showed a lower percentage of MYC footprints localizing downstream of TSSs compared to sites that co-bound MYC and RNAPII (46.0% +/- 1.7% vs 54.1% +/- 1.1%, p=0.015) but equal amounts of MYC binding downstream of TESs (59.1% +/- 2.0% vs. 60.0% +/- 3.0%, p=0.40) (Supplementary Tables 3 and 6). The former observation likely reflected MYC’s absence at downstream sites of transcriptional pausing (Figure 1A) (8). Differences between the above two groups included the already noted greater amount of TSS- and TES-associated MYC binding in transformed versus non-transformed cell lines (18–21, 27). Although MYC-binding TESs accounted for 10.2% of all gene-associated binding, it was over five times more common in RNAPII’s absence than in its presence. This lower overall frequency of TES-associated MYC binding relative to the previously reported ∼17.5% can be attributed to the fact that the latter also included sites occupied by MAX-MXD heterodimers (27).

Genes associated with only RNAPII at TSSs and/or TESs showed levels of binding downstream of their respective sites very similar to those seen when it co-bound with MYC (62.7% +/- 1.4% vs. 62.8% +/- 0.9% for TSSs [p=0.97] and 60.3% +/- 1.1% vs. 58.9% +/- 1.8% for TESs [p=0.58]) (Figure 1E and F and Supplementary Tables 3 and 7). Collectively, these results demonstrate that the fractional binding of MYC and RNAPII with respect to TSSs and TESs is significant only for case where the greater amount of MYC binding downstream of MYC+RNAPII co-bound TSSs may reflect MYC’s predilection to associate with RNAPII at transcriptional pause sites (8).

### MYC and RNAPII show distinct sequence-specific binding preferences at TSS- and TES-proximal sites that are also enriched for other TF binding motifs

Genes identified as bearing MYC and/or RNAPII ChIP footprints at TSS and/or TESs were arranged into eight alphabetical and 16 numerical categories that accounted for all possible binding combinations (Figure 2A, 2B, Supplementary Figure 1, Supplementary Files 1 and 2). The former (peak assignments) represented each of the possible MYC/RNAPII binding configurations at either TSSs or TESs without regard to the binding status of the other end of the gene, whereas numerical categories (gene assignments) took this into consideration. “Analysis of Motif Enrichment” (AME) and “Find Individual Motif Occurrences” (FIMO) tools were then used to identify consensus TF binding sites localizing to within +/-50 bps of all MYC and RNAPII peak summits (51). From among the 879 consensus motifs in the JASPAR database (https://jaspar.elixir.no/), 559 (63.6%) were significantly enriched within these short DNA segments in at least one of the above categories (52). This showed that each alphabetical category could be distinguished based solely upon the content of its associated TF binding sites (Figure 2C, 2D, and Supplementary Files 3-10).

**Figure 2.**
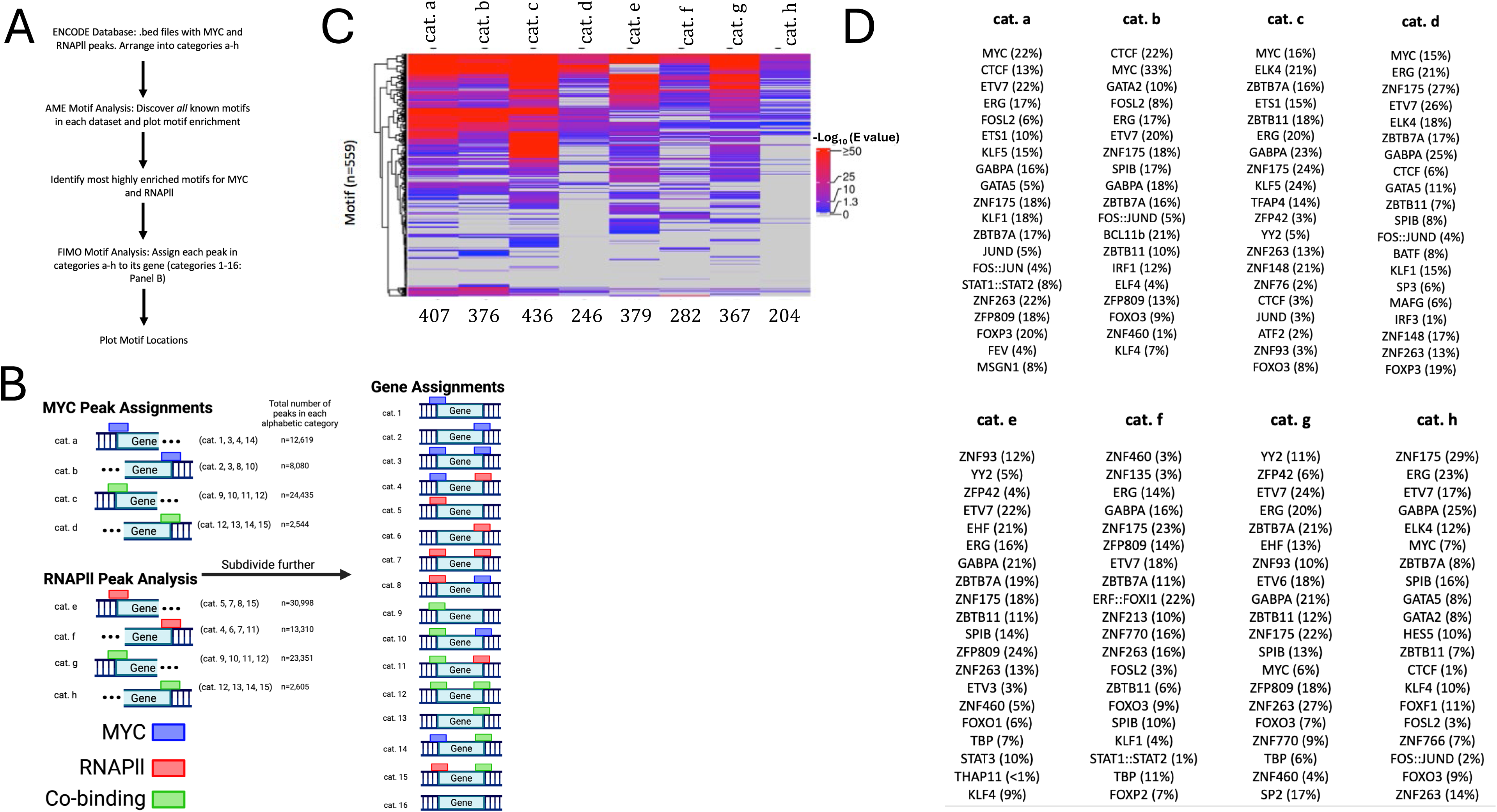
Association of MYC and/or RNAPII with consensus DNA binding site motifs. (A). Work flow diagram for identifying MYC and RNAPII ChIPseq peaks and their associated DNA binding motifs at TSSs and TESs. The “Assessing Motifs Enrichment” (AME) tool was first used to identify putative TF binding motifs associated with regions +/- 50 bps flanking the ChIPseq peak summits from the ENCODE database for all cell lines. The motifs identified were then further refined and more precisely mapped within their respective genes using the “Find Individual Motif Occurrences” (FIMO) tool, which has stricter identification criteria than AME. For both analyses, shuffled primary sequences were used as an internal control (51). (B). Alphabetical (peak assignment) and numerical (transcript assignment) categories based upon MYC and RNAPII binding site combinations around TSSs and TESs as described in (A). The majority of MYC and RNAPII ChIPseq peaks were mapped to within +/-500 bps of TSSs and to within +/- 2.5 kb of TESs as previously described for MYC and MAX (Figure 1) (27). The dotted lines associated with alphabetical categories a-h indicate that these assignments were made without regard to the binding of MYC and RNAPII at the other ends of their individual genes. Co-binding sites were defined as MYC and RNAPII peaks sharing ≥100 bp of overlap (91). Note that categories c & g and d & h, while appearing identical in the cartoon, were based on ChIPseq experiments performed with different antibodies (MYC for categories c and d and RNAPII for categories g and h). The locations of these peak summits and their associated TF binding motif subsets thus typically differed. See Supplementary Figure 1 for a more detailed presentation of how categories a-h were identified and assigned. For subsequent gene expression analyses, these eight alphabetical categories were subdivided into 16 numerical ones (cats. 1-16) based on whether MYC and/or RNAPII binding were associated with TSSs and/or TESs as previously done for MYC and MAX binding (27). Numbers to the right of each alphabetical category indicate the total number of genes bearing the indicated MYC/RNAPII configurations for all seven cell lines. See Supplementary File 1 for the raw ChIPseq Peak sequences associated with each alphabetical category and Supplementary File 2 for the transcript IDs associated with each numerical category. (C). Heat maps of TF-binding motifs residing within +/- 50 bps of MYC and RNAPII ChIPseq peak summits at TSSs and TESs of gene categories a-h from panel B. The 559 motifs shown were derived from a total of 879 in the JASPAR data base. Results from all seven evaluable cell lines from the ENCODE database have been combined and are shown here. Rankings were based on E values (52). Numbers beneath each column indicate the number of statistically significant enriched motifs identified in the combined cell-line data. Note that an enriched motif in this plot does not guarantee its enrichment in any individual cell line, although it will correlate. (D). Rank lists of the ∼20 most significantly associated DNA binding motifs from panel C. All results shown have E<0.05. See Supplementary Files 3-10 for individual cell line results. The consensus motifs for MYC, MYCN, MAX, ARNT, CLOCK and USF1 motifs, each of which was separately identified as being among the top 20 of all sites (Supplementary Figure 2), have been combined here and labeled as “MYC” sites as they all contain similar CACGTG core elements capable of binding MYC-MAX heterodimers with varying affinities (53, 54, 92). SP1 sites, while significantly enriched, were not always identified as among the top 20 DNA motifs but are included nonetheless due to their well-known ability to interact with MYC-MAX heterodimers and suppress gene expression (14, 16, 92). That the percent of motifs shown exceeds 100% is consistent with the fact that footprints were typically associated with multiple TF binding motifs as noted in (C) and elsewhere (27). Note that cat. b only has 19 entries as there was only 19 significantly enriched and distinct TF motifs.

Unsurprisingly, among the top motifs identified with MYC ChIPseq peak summits, regardless of whether they localized to TSS- or TES-proximal sites or were associated with coincident RNAPII binding, were canonical E boxes (CACGTG). Because these often differed in their flanking sequences, they were sometimes classified in the JASPAR database as being preferred sites for bHLH-ZIP or bHLH-PAS TFs other than MYC such as MYCN, CLOCK, ARNT and USF. These motifs may nonetheless still serve as *bona fide,* albeit sometimes less avid, MYC binding sites, particularly in cells with high MYC levels (27, 53–57). We therefore initially classified them as generic MYC E box elements (Figure 2D). Consensus binding motifs for CTCF and members of the FOS-JUN family of bZIP TFs were also enriched but were at least 3-4 times more common at TSS- and TES-proximal sites that bound only MYC (cats. a and b). The close proximity of these to MYC peak summits can likely be explained by the fact that CTCF directly interacts with MYC whereas FOS-JUN heterodimers and JUN homodimers often recognize consensus AP1 sites that reside in close proximity to E boxes (27, 58–61). Canonical SP1/3 sites, while not among the top 20 enriched motifs in all alphabetic categories, were nonetheless still significantly enriched beneath MYC ChIPseq TSS and TES peak summits, as well as beneath RNAPll ChIPseq peak summits at TSSs (E < 0.05). On the other hand, KLF5 motifs, some of which also bind SP1 (62) were identified in association with MYC and MYC+RNAPII binding TSSs in 15% and 24%, respectively of cats. a and c genes but not in any other alphabetical categories. KL4 motifs, which are virtually identical to SP1 sites, were seen at much lower abundance (7-10% of sites) in genes comprising cats. b, e and h. ZBTB17 motifs were not highly enriched among any categories. Thus, from among the top ∼20 consensus TF binding motifs most strongly associated with MYC ChIP footprints, canonical E boxes, or sites that bind MYC-interacting TFs, were found to reside within +/- 50 bps of many peak summits. While the remaining motifs almost certainly do not bind MYC directly, their non-random clustering in extreme proximity to MYC binding sites suggested that they may associate with their cognate TFs in ways that indirectly influence MYC-mediated transcription and *vice versa* (27, 33, 41, 63, 64).

Breakdown of the above generic E box motifs into more specific subsets based on their flanking sequences showed significant differences in their association with MYC ChIP peak summits at TSS- and TES-proximal sites (Supplementary Figure 2A and B). As expected, there was also more enrichment of sites associated with MYC ChIP peaks (cats. a-d) than with RNAPII ChIP peaks (cats. e-h). Finally, there was a ∼50% higher level of enrichment of MYC and MAX-specific sites at TSSs and TESs occupied only by MYC (cats. a and b) versus MYC+RNAPII (cats. c and d).

In contrast to the MYC ChIPseq results, the motifs most frequently associated with RNAPII binding peak summits were more diverse and less likely to be enriched for E boxes and SP1/KLF sites (Figure 2D [cats. e-h] and Supplementary Files 3-10). The latter motifs, specifically, those labeled as KLF4-like were identified at TSS-associated RNAPII only sites (cat. e: 9%) and at TES-associated MYC+RNAPII co-binding sites (cat. h: 10%). The frequency of some motifs also differed at TSSs and TESs. For example, YY2 and ZNF93 binding sites were associated with 5-12% of TSS-associated peak summits but were not among the top motifs associated with TES-associated peak summits. In contrast, motifs for other TFs, including GABPA, ZNF175, and ERG, were identified with similar frequencies across all four categories, whereas other consensus motifs such as those for FOXI1, HESS and ELK4 were associated with only a single binding category (e.g. cats. f or h).

Even within the tight +/-50 bp confines of the MYC footprint peak summits displayed in cats. a-d, many of the above-mentioned TF binding motifs were not distributed randomly. For example, E boxes were always found in closest average proximity to MYC peak summits, regardless of whether they resided at TSSs or TESs or co-localized with RNAPII (Figure 3A). This precision provided additional strong evidence for actual direct binding of MYC-MAX heterodimers to these sites. Other motifs, such as those for ETS1 in cat. a, ERG in cat. b, and GABPA in cat. d were also non-random in their distribution but tended to localize at sites more remote from MYC peak summits. In contrast, many of the remaining TF motifs such as those for ZFP809 in cat. a, GATA2 in cat. b, FOXO3 in cat. c and BATF in cat. d were distributed more randomly. Together, these findings provided further evidence that at least some of the numerous other TF binding sites reside at recurrent and precise distances from MYC peak summits and that their positioning is not identical between TSSs and TESs.

**Figure 3.**
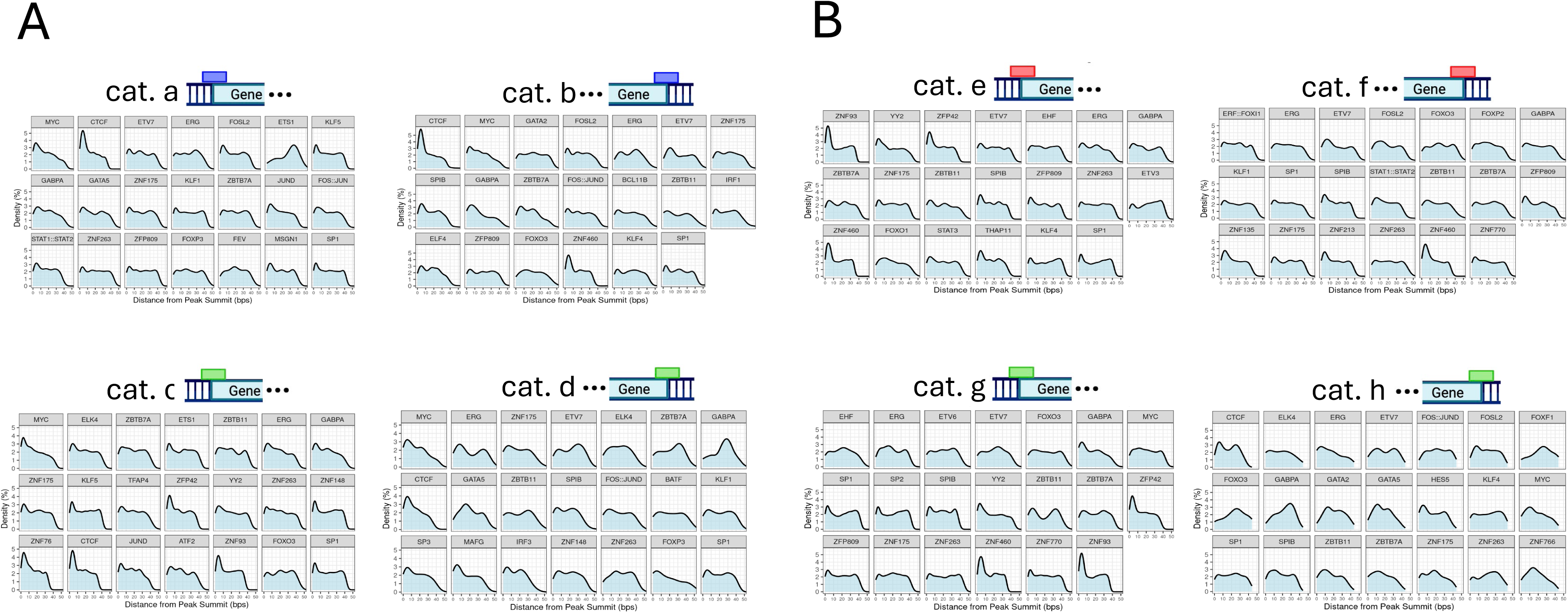
Distribution of TF binding motifs around MYC and RNAPII ChiP peaks from all seven cell lines. Motifs listed in Figure 2D were subject to FIMO analysis with the output peaks being mapped back to the original ChIPSeq peaks. This allowed for an estimate of the distance from the beginning of each motif to its respective peak summit. Aggregate distances were then plotted for each motif as a distribution. (A). Distribution of the average distance from the end of a consensus DNA binding motif to its peak summit summits in alphabetical categories a-d. (B). Distribution of the average distance from the end of a consensus DNA binding motif to its peak summit summits in alphabetical categories e-h.

Similar findings were made with respect to RNAPII peak summits at TSSs and TESs (cats. e and f, respectively) (Figure 3B). For example, ZNF460 motifs were among those residing in closest proximity to the summits in both categories. In contrast, other motifs, such as those for ZBTB11 in cat. g and FOXF1 in cat. h clustered at more remote sites. As previously seen with MYC binding, many motifs were also distributed relatively randomly with respect to peak summits and included such sites as ZNF263 (cat. e), ERG (cat. f), FOXO3 (cat. g) and FOS:JUND (cat. h). Notably, MYC motifs associated with cat. g genes were more randomly distributed around RNAPII peak summits indicating that these sites, while close by, were unlikely to be primary sites of RNAPII binding (53).

An interesting finding that applied to both MYC and RNAPII was that despite the diversity of enriched TF binding sites in proximity to TSSs and TESs, more than half were not only common but often clustered similarly around peak summits. In the case of MYC for example (cats a and b), relevant motifs included those for MYC, CTCF, FOSL2 and GABPA. Similarly, in the case of RNAPII (cats. e and f), these included other consensus sites for MYC and CTCF as well as those for unique factors such as ETV7, ERG, and ZNF175. Along with the absence of MYC and/or RNAPII at some of these sites, these findings implied that the binding of both MYC and RNAPII was a primary event rather than a passive one due to processive transcription initiated at promoters. The abundance of E boxes at TESs that were otherwise occupied only by RNAPII also suggested that such sites might be occupied under different circumstances (see below).

### MYC and RNAPII binding around TSSs and TESs correlate with gene expression in dynamic and reversible ways

The expression of genes associated with each numerical category (Figure 2B) was next quantified, with particular focus on variations among subsets with MYC and/or RNAPII binding at different sites. For all seven cell lines, we compiled ENCODE database RNAseq results and compared mean absolute TPM levels for each category. Combining the results increased the power of these comparisons while minimizing any cell type-specific or other idiosyncratic contributions (Figure 4 and Supplementary Table 8). As expected, cat. 16 genes (devoid of any MYC or RNAPII binding) were expressed at the lowest average level (27) (Figure 4A). In contrast, the mean expression levels of cats. 1, 5, and 9 genes, which bound MYC, RNAPII or both factors only at TSS-proximal sites were 3.4-, 5.2- and 7.9-fold higher, respectively, than cat. 16. In comparison, the binding of MYC, RNAPII or both factors at TES-proximal regions only (cats. 2, 6, and 13) was associated with 2.2, 9.2 and 3.0-fold higher levels of expression, respectively (Figure 4B). This indicated that, as a group, genes with RNAPII binding only at TESs (cat. 6) were expressed at higher levels than those of both cat. 16 and cat. 5, which bound RNAPII only at TSSs. It further indicated that MYC binding at TESs, either alone or with RNAPII, was associated with a lower average gene expression level when adjusted for their binding at TSSs.

**Figure 4.**
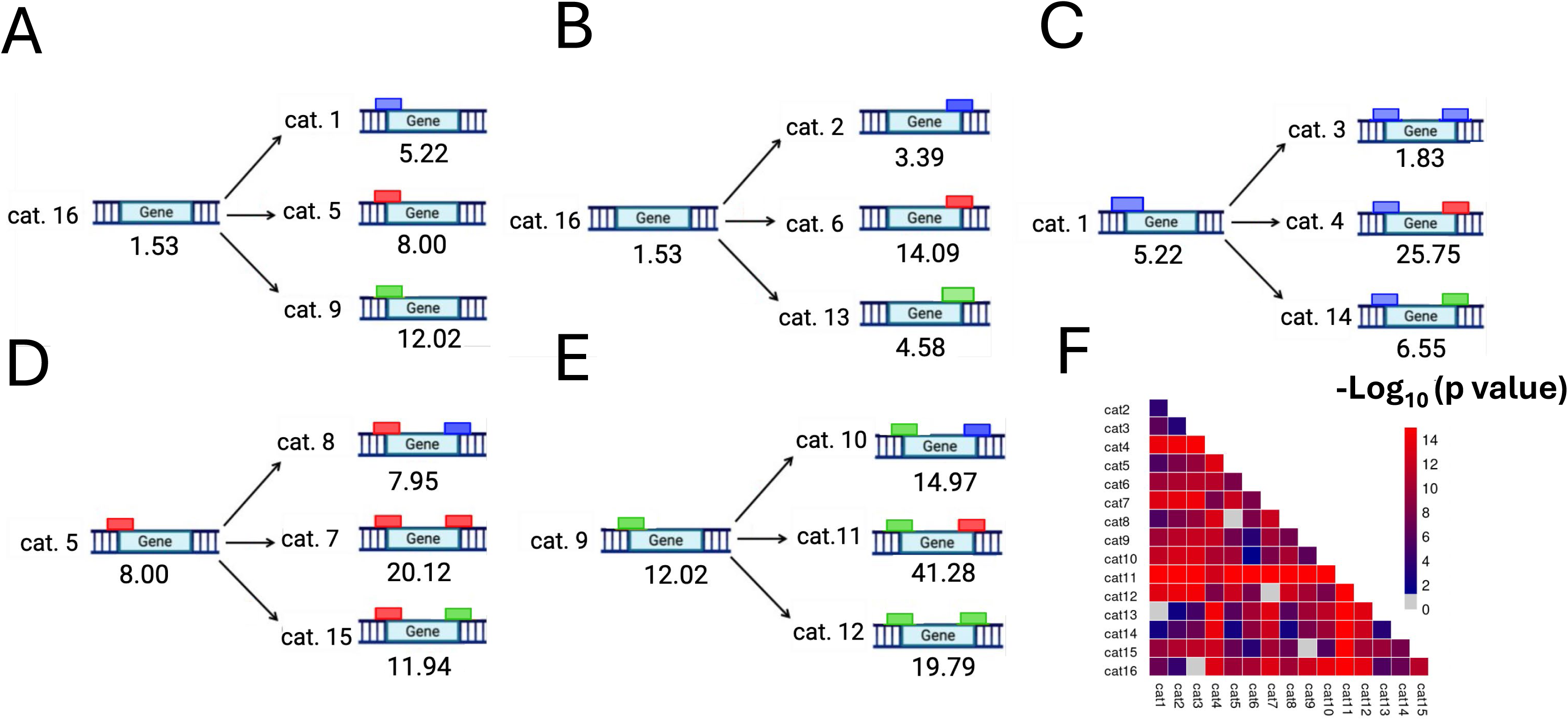
Mean transcript levels of genes in the indicated numerical categories. (A-E). Raw TPM values were extracted from RNAseq data without further pre-processing. Transcript IDs from each numerical category were matched to those in the RNAseq data and non-matching transcripts (<10%) were eliminated. Aggregate averages for each category and cell line were calculated, with each cell line’s values being normalized to its own mean to account for inter-cell line differences. Normalized cell-line means were further combined to create a single value for each category describing the transcript level of genes in that numerical group. Listed below each of the indicted gene categories are their mean raw TPM values across all cell lines. See Supplementary Table 8 for expression values for individual cell lines. (A,B). Influence of MYC and/or RNAPII binding at TSSs and TESs, respectively, on the expression levels of the indicated categories of genes relative to those of cat. 16. (C). Relative effect of MYC and/or RNAPII binding in proximity to TESs in genes that bind only MYC around TSSs. (D). Relative effect of MYC and/or RNAPII binding in proximity to TESs in genes that bind only RNAPII around TSSs. (E). Relative effect of MYC and/or RNAPII binding in proximity to TESs in genes that co-bind MYC and RNAPII around TSSs. (F). Significance of pair-wise differences in mean gene expression levels among each of the indicate gene categories. Each value was calculated via a paired t-test. p-values from each result were transformed via - log_10_ and plotted as a heatmap.

Somewhat different behaviors were observed among the gene categories that, in addition to binding MYC, RNAPII or both factors at TESs, also bound MYC at TSSs (Figure 4C). For example, and as already noted, cat. 1 genes were, on average, 3.4-fold more highly expressed than cat. 16 genes. Genes with additional TES-proximal binding of MYC (cat. 3), RNAPII (cat. 4) or both factors (cat. 14) showed either a 65% lower mean level of expression (relative to cat. 1) or 4.9 and 1.3-fold higher levels, respectively. The 3.9-fold lower level expression of cat. 14 relative to cat. 4 indicated that, in this context, MYC binding at TESs was again relatively suppressive.

Similar findings were made with respect to cat. 5 genes, which bound only RNAPII at TSSs and were associated with 5.2-fold higher mean expression levels relative to cat. 16 genes (Figure 4D). Additional MYC binding at TESs (cat. 8) was associated with no change in mean expression whereas TES-associated RNAPII binding genes (cat. 7) were expressed at 2.5-fold higher levels. Once again, MYC+RNAPII at TESs (cat. 15) marked genes whose expression was suppressed relative to those binding only RNAPII (cat. 7).

Finally, the above-noted trends were largely mimicked among genes that bound MYC+RNAPII at TSS-proximal sites (cat. 9) (Figure 4E). This alone was associated with a 7.9-fold higher average mean expression level versus cat. 16. Additional MYC binding at TESs (cat. 10) was associated with a further modest increase in average gene expression (1.2-fold), whereas RNAPII binding in genes (cat. 11) was associated with 3.4-fold higher level of expression. As before, genes with dual binding around TESs (cat. 12) were expressed at lower levels (1.6-fold higher than cat. 9), which again pointed to the repressive nature of MYC at TESs in association with RNAPII. Comparisons between all pairwise groups were visualized with a p-value heatmap (Figure 4F).

Many MYC- and/or MAX-bound genes undergo “category switching”, particularly when MYC levels are changing rapidly (27). To examine this in the context of MYC and RNAPII binding and whether it correlated with alterations in gene expression, we retrieved ChIPseq and RNAseq data sets from murine livers, which express extremely low levels of endogenous Myc, and compared the findings to hepatocellular carcinomas (HCCs) that rapidly arise in response to the conditional, high-level induction of a human *MYC* transgene (65, 66). We first identified a large group of genes in livers that bound only RNAPII at TSSs and were thus representative of cat. 5 (Figure 5A). In HCCs, all these genes switched to cat. 9 as a consequence of additional MYC binding near these same sites (cat.5→cat.9 switching). Many, if not all, of these genes thus likely contain either low-affinity MYC binding sites in their promoters and/or higher affinity sites that were initially inaccessible to MYC (27).

**Figure 5.**
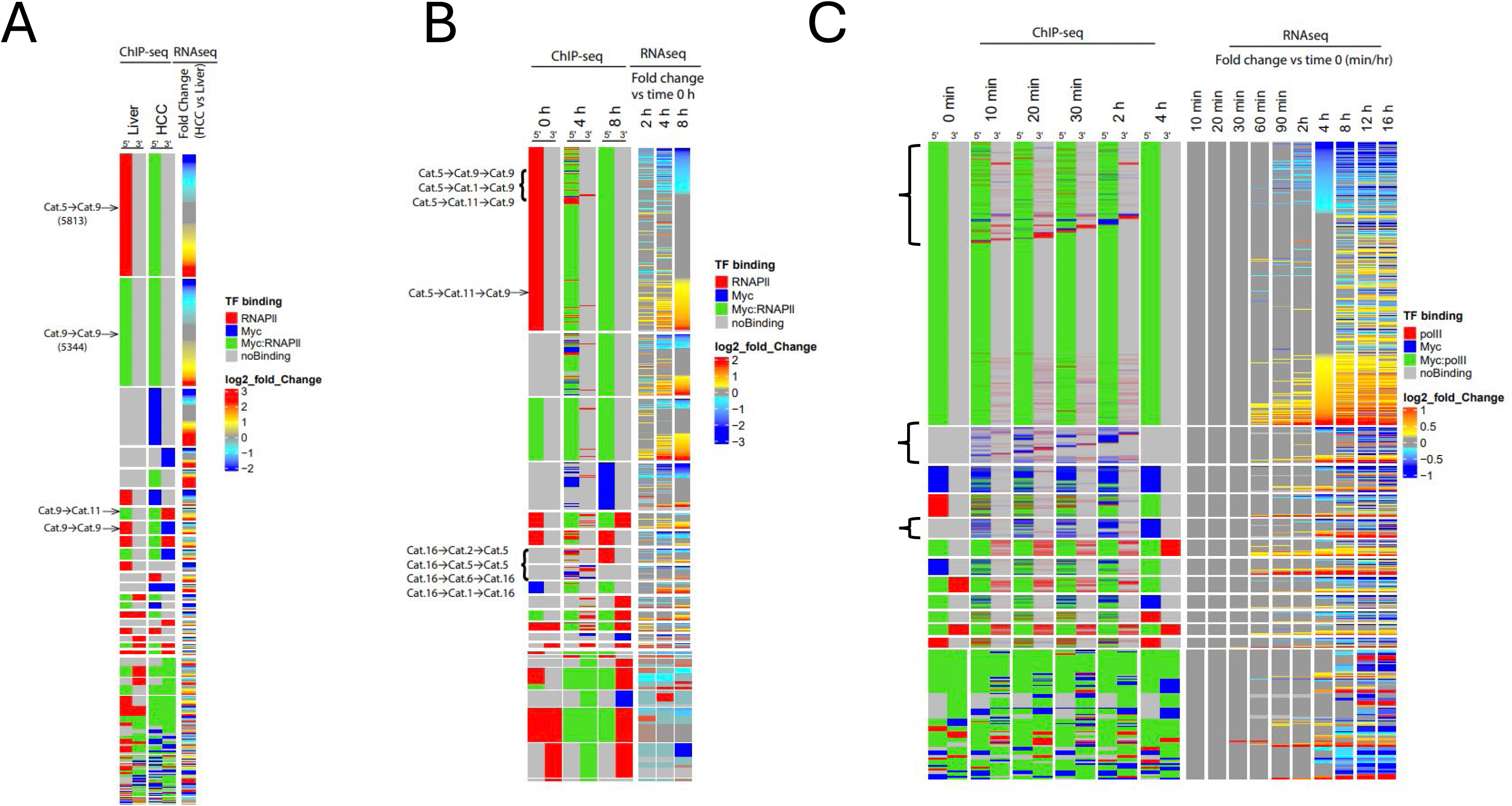
MYC and RNAPII binding at TSSs and TESs is dynamic and reversible in response to oncogenic and mitogenic signaling. (A). ChIPseq and RNAseq results from control livers and MYC-induced HCCs (65, 66). Numerical gene categories, identified by ChIPseq in control murine livers are indicated in the first two columns, and those from HCCs are indicated in the third and fourth columns. The designations 5’ and 3’ represent binding by MYC and/or RNAPII to TSS- and TES-associated sites, respectively. In this way, category switching can be readily identified, with relevant examples being indicated by black arrows. The last column summarizes RNA-seq results, with log_2_ fold change values shown on the indicated color scale. Blue and red denote genes that are down- and up-regulated, respectively, in HCCs versus livers. (B). ChIPseq and RNAseq results from LPS-treated murine B cells (27, 67). Columns are designated as described in (A). Three sets of columns are displayed representing ChIPseq results from control, quiescent B cells (0 hr) and those treated with LPS for four and eight hr. The rightmost three columns show gene expression profiles obtained two, four and eight hr after LPS treatment with the levels of genes being compared to those in untreated B cells (0 hr). (C). ChIPseq and RNAseq results from serum-treated murine 3T9 fibroblasts (27, 68). In addition to the first two columns, representing cells maintained under low serum conditions, ChIPseq and RNAseq results from five time points following serum stimulation are shown. Note that the most abundant group of genes are those classified as cat. 9 prior to and following a 4 hr. exposure to serum. Within this group however are numerous genes that show dynamic category switching at intermediate times following serum stimulation (upper bracket). The lower two, smaller brackets shows examples of cat.16 genes at time 0 that undergo several different types of transient changes, eventually returning to their original cat. 16 status or transitioning into cat. 1 genes.

A second, equally large group of liver genes that bound MYC+RNAPII only at TSSs remained unchanged in HCCs (cat. 9→cat. 9) (Figure 5A). The prominent MYC ChIP profiles associated with these sites in livers with barely detectable levels of MYC expression thus indicated that they were of high-affinity and thus distinct from the previously-mentioned cat. 5 genes. These findings also indicated that many genes are not necessarily associated with category switching following neoplastic conversion. Other notable groups were comprised of smaller gene sets that showed no evidence of TES-associated MYC or RNAPII binding in livers but subsequently acquired these profiles in tumors. These included cat.16→cat.2, cat. 9→cat.11 and cat. 5→cat.10, with the latter showing switching at both TSS- and TES-associated sites. Collectively, these findings indicated that, like much of their binding at promoters and enhancers, MYC and RNAPII binding at TES-associated sites can involve diverse types of category switches. The paucity of genes with TES-associated MYC binding in control livers with extremely low levels of endogenous MYC also confirmed our previous findings that these sites are, on average, of lower affinity and/or are otherwise inaccessible (27). Virtually all of the categories ultimately associated with HCCs contained gene subsets whose expression was increased, decreased or unchanged relative to livers (Figure 5A).

Because the above results were generated from only two points (i.e. livers and HCCs), they were unable to address such questions as whether category switching proceeds at variable rates, whether multiple category switches in single genes can occur and whether category switching is reversible. To answer these, we relied upon two additional ChIPseq/RNAseq datasets with multiple time points. The first, generated from lipopolysaccharide (LPS)-treated B cells (27, 67), allowed us to draw several conclusions. First, many of the same numerical categories described in livers were also identified in quiescent B cells although the identities of the component genes differed (Figure 5B). Most of the categories contained gene subsets that underwent at least two distinct category switches during the course of LPS treatment. Category switches often occurred at different rates and some were reversible despite the continuous presence of LPS and ongoing cell cycle progression. In other cases, switching from one category to another proceeded along different routes, for example via cat.5→cat.9 directly or via intermediates such as cat.5◊cat.1◊cat.9 or cat.5◊cat.11◊cat.9. The generality of these findings was confirmed in a third ChIPseq/RNAseq data set from serum-treated NIH3T9 fibroblasts (68). The larger number of time points obtained over a four hr. period of sampling again showed ongoing multiple category switches, with some recapitulating previously observed categories (Figure 5C, brackets). A particularly notable group of genes were those with a cat. 9 binding pattern at 0 and 4 hr that nonetheless showed multiple patterns of category switching at intermediate times (10 min-2 hr). This indicated that, at least in serum-stimulated fibroblasts, much of the TES-associated MYC and RNAPII binding is observed only during the brief transition from a state of quiescence to one of active proliferation. As was true for MYC-driven HCCs, most categories from these mitotically-stimulated B cells and fibroblast cells contained gene subsets whose expression increased, decreased or remained unchanged relative to their unstimulated counterparts.

### Numerical categories contain functionally-enriched gene sets

Given the extensive variability of inter-category switching in response to normal and neoplastic mitogenic signals (Figure 5), we were interested in determining whether individual categories contain functionally related genes at times when MYC levels remained constant. We suspected that this might allow these genes to be coordinately regulated in response to altered demands for MYC-regulated activities such as ribosomal biogenesis, mRNA splicing, and cell cycle progression (2, 6, 11, 17, 65). This notion was supported by our previous work indicating that genes with TES-associated MYC and/or MAX binding play disproportionate roles in proliferation, differentiation and various immune and inflammatory responses (17). We identified 5699 gene sets from the Organism Database (OrgDb) that were significantly enriched in at least one category in one or more cell lines (Figure 6A and Supplementary File 11). Among the findings revealed by this analysis was that the two cell lines with the highest levels of gene set enrichment (MEL and CH12.lx) were those with the highest MYC levels (Supplementary Table 1). These patterns were generally not maintained across all cell lines or, if they were, were enriched to variable degrees. This suggested, as previously noted above and elsewhere, that the identities of genes within each numerical category were not only sensitive to MYC levels but were cell type-specific as well (Figure 5) (11, 17).

**Figure 6.**
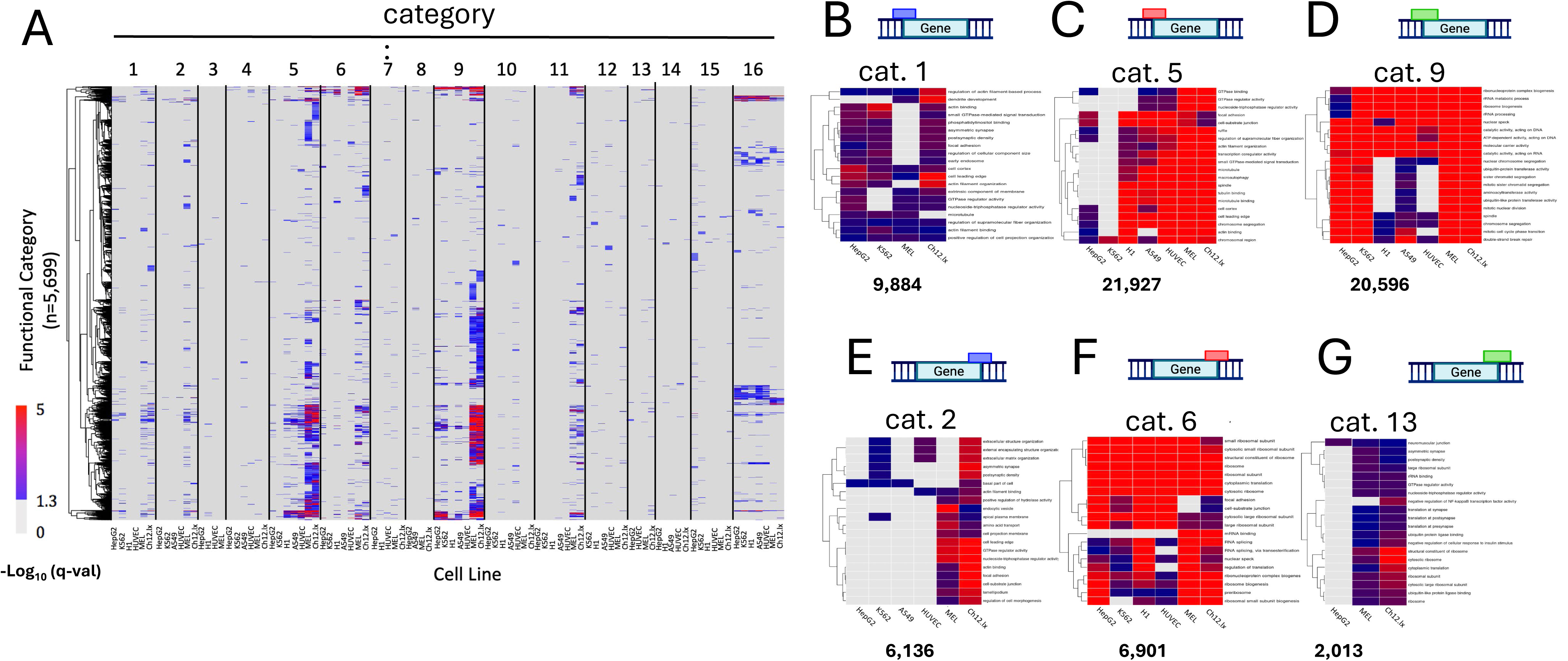
Different numerical gene categories are associated with specific functions. (A). For each of the seven cell lines, transcripts from each numerical category were compared to gene sets from the Organism Database (OrgDb: https://orgdb.github.io/) to determine gene set enrichment. Each tick on the subsequently constructed heat map indicates one of the 5699 gene sets that was significantly enriched in at least one cell line. The absence of some cell lines in certain categories shown in the panel (e.g. cat. 3 and cat. 8) indicates that no gene sets were significantly enriched and/or that too few genes comprised the category to allow for meaningful screening. All three ontologies in the OrdDb database were retained for this analysis. (B-G). The top 20 most highly enriched gene sets from among those shown in A are depicted to allow for shared functions across tissues and species to be better appreciated. Numbers beneath each panel indicate the total number of binding sites for the indicated factors across all cell lines. See Supplementary File 11 for a complete list of all enriched gene sets.

Cats. 1, 5, and 9, whose members lacked any TES-associated MYC or RNAPII binding, contained the largest total number of genes, with MYC, RNAPII or MYC+RNAPII being associated with between 9884-21927 TSS-proximal binding sites (Supplementary Table 9). They also contained the largest number of enriched OrgDB gene sets (Figure 6A). Although many of these were cell line-specific, others were shared across as many as all seven cell lines. For example, among the top 20 most heavily enriched cat.1 gene sets were four whose members participated in actin organization or function in three or four different cell types (Figure 6B). The top 20 gene sets in cat. 5 contained some of the same enriched sets while including others related to GTPase binding and activity and mitotic spindle/tubulin and chromosome structure and function (Figure 6C). Finally, over half of the top 20 enriched gene sets for cat. 9 were related to ribonucleoprotein and rRNA biogenesis as well as chromosome/chromatid segregation, spindle formation and mitosis (Figure 6D). Twelve of these top 20 gene sets were enriched in all seven cell lines and perhaps reflecting a combination of non-mutually exclusive factors such as their fundamental importance, their content of high affinity MYC binding sites, and the independence of MYC and RNAPII binding on tissue-specific co-factors (69, 70).

Cats. 2,6 and 13, whose member genes bound MYC, RNAPII or both factors only at TES-proximal sites, also contained significantly enriched gene sets although these tended to be more diverse and cell line-restricted (Figure 6E-G and Supplementary File 11). For example, among the most highly-enriched gene sets for cat. 2, only six were enriched in three cell lines and only one was enriched in four (Figure 6E). Similarly, among the 20 most enriched gene sets for cat. 13, only one was enriched in three cell lines with the other 19 gene sets being similarly enriched only in MEL and CH12.lx cells with the highest levels of MYC, and again supporting the idea that MYC binding around TESs is of lower average affinity (Figure 6G) (27). Cat. 6 was a marked exception to the others as, in the six evaluable lines, 14 of the 20 most significantly enriched gene sets involved functions pertaining to translation or ribosome structure and function, with 12 of these being enriched across all cell lines. Moreover, of the remaining six enriched sets, four involved functions that are closely linked to translation (i.e. splicing, mRNA binding and nuclear specks). Collectively, these results indicate that, at least under conditions where MYC levels remain constant, genes with specific patterns of TSS- and TES-associated MYC and/or RNAPII binding sites tend to identify distinct tissue-specific and shared functional genes groups. The most striking of these are evidenced by cat. 6 genes, which are heavily enriched for the well-known MYC-regulated functions of ribosomal biogenesis and translation (6, 11, 17, 71).

### Read-through transcription is jointly regulated by TES-proximal MYC and RNAPII

Read-through or so-called “downstream of gene” (DoG) transcription describes the passage of RNAPII beyond the normal, 3’-most termination/polyadenylation site, sometimes by as much as 50-100 kb (31, 32). It occurs in selective genes in a wide range of normal and neoplastic tissues and often increases significantly in response to stresses such as heat, hypoxia and oxidative or osmotic shock (27, 28, 31, 32). We have previously demonstrated that genes with TES-associated MYC binding often display transcriptionally conducive histone modifications around these sites, an open local chromatin environment, and higher levels of E box-dependent DoG transcription following exposure to some of these stresses (27). However, the variability of the responses suggested that MYC’s participation in DoG transcription requires other factors and/or that MYC’s role is stress type-dependent. We thus examined the degree to which DoG transcription of protein-coding genes in the previously studied cell lines was also influenced by TES-proximal RNAPII binding, either independent of or in association with concurrent MYC binding. For this, we combined RNAseq results for each cell line from numerical cats. 2,3,8, and 10 (MYC only binding at TESs); cats. 4,6,7 and 11 (RNAPII only binding at TESs); cats. 12-15 (MYC+RNAPII co-binding at TESs); and cats. 1,5, and 9 (neither MYC nor RNAPII binding at TESs) (Figure 2B). The fractional amount of DoG transcription for each gene in these categorical groups was then determined by adjusting this value to that of total gene expression (27). Although significant differences were sometimes observed among each of these groups, they were almost always small and highly tissue-specific, with neither MYC nor RNAPII being reproducibly associated with either higher or lower levels of DoG transcription.

Our demonstration that specific patterns of TES-proximal MYC and/or RNAPII binding identified categories of genes with related functions did not initially consider their DoG transcription status (Figure 6). When these results were re-analyzed to account for this, genes associated with DoG transcription were found to contain very different GSEA profiles than those without (Figure 7B and Supplementary Files 12 and 13). Most strikingly, and in both human and mouse cells, functional enrichment of genes that bound MYC and/or RNAPII at TES-associated sites was most pronounced for those that were DoG transcription +. Re-examination of the 20 most significantly enriched gene sets from the four groups further demonstrated the extent to which certain functional groups were marked in this manner. For example, DoG transcription + gene sets associated with MYC only TES binding were notable for their roles in regulating protein localization and GTPase binding in human cells and, in murine cells, ribosomal and mitochondrial processes (Figure 7C). The top DoG transcription + gene sets associated with RNAPII binding only at TESs in both human and murine cells were those with roles in ribosome rRNA structure and function, splicing, and ribosomal/RNA-protein interactions (Figure 7D). In the MYC+RNAPII co-binding category, cytoskeletal-related gene sets were most enriched in human cells while neuron-specific translation genes were enriched in murine cells (Figure 7E). Finally, human gene sets with neither MYC nor RNAPII at TES-proximal sites were particularly enriched for those encoding proteins with roles in membrane functions and phosphate metabolism; murine cells had similar enrichment of membrane-related genes and species-specific ones relating to cell polarity (Figure 7F). Collectively, these results indicate that MYC and/or RNAPII binding at TES-proximal sites tends to selectively mark DoG + transcription subsets of genes that share common functions.

**Figure 7.**
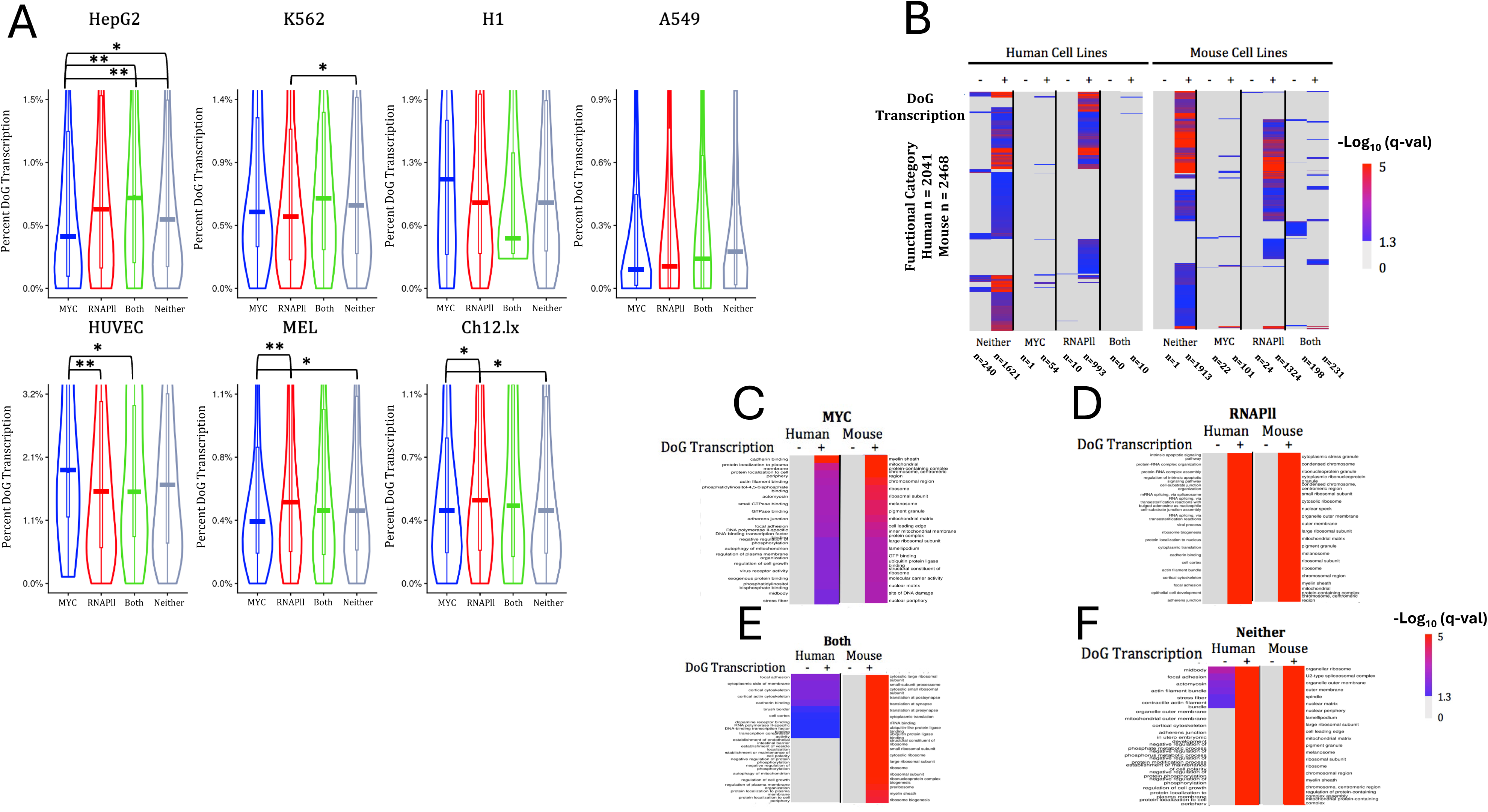
MYC and/or RNAPII binding at TES-proximal sites marks genes with higher levels of DoG transcription and distinct functions. (A). Fractional DoG + transcription for expressed coding genes in the indicated cell lines. As described in Figure 6, genes from all seven cell lines were combined and placed into four groups based on whether they bound MYC only (numerical cats. 2,3,8, and 10 [Fig. 2B]); RNAPII only (cats. 4,6,7, and 11), MYC+RNAPII (cats. 12-15), or neither factor (cats. 1, 5, and 9) at TES-proximal sites. In the case of genes with multiple TESs, only the most 3’-TES was used to identify DoG transcription so as to avoid the inclusion of genes that were merely utilizing alternative cleavage sites. n = the total number of genes in each of the indicated groups. Numbers in parentheses indicate the percent of these showing evidence of DoG transcription. *: p<0.10; **: p<0.05. Comparisons were performed using the Mann-Whitney U-test. See Supplementary File 12 for a compiled list of DoG readthrough values. (B). MYC- and/or RNAPII-associated DoG + expression identifies distinct functional categories of genes in both human and murine cell lines. Gene groups were the same as those described in A. They were then further divided according to whether or not they displayed evidence of DoG transcription. Finally, GSEA was performed upon the DoG+ and DoG- groups with the 5999 gene sets from the OrgDb to identify functionally enriched gene sets. Numbers along the vertical axis of each panel indicate the total number of gene sets that were significantly enriched in at least one of the four groups. Numbers beneath each column indicate the number of genes for which DoG transcription was or was not detected. See Supplementary File 13 for a list of compiled significantly enriched gene sets. (C-F). Top 20 most significantly enriched gene sets from the DoG+ and DoG- groups in B. Again, see Supplementary File 13 for a comprehensive listing of all significantly enriched gene sets from each of these groups.

## Discussion

The binding of MYC and/or MAX to TES-proximal sites in approximately one-sixth of all annotated genes occurs in different contexts and exerts variable qualitative and quantitative effects on gene expression (Figure 4A-C, Figure 5) (27). Unlike the 5’-ends of genes where 90% or more of all detectable MYC footprints map to within 1 kb of TSSs, only about half of the footprints associated with 3’-ends cluster this closely to TESs while also showing somewhat lower overall mean binding affinities (Supplementary Table 2) (27). Despite these differences, TSS- and TES-associated MYC-MAX and MXD-MAX binding share several important properties. Chief among these is that much of the binding occurs directly at E boxes or indirectly at SP1/3, ZBTB17, and CTCF sites which often bind their eponymous TFs in association with MYC-MAX heterodimers (Figure 2D and Supplementary Files 3-10) (13–18, 27, 41, 53, 60, 66). Second, many of these sites are of low affinity and are occupied only in certain cancers with high MYC levels (27, 65, 66). Third, the order in which genes with multiple TSS- and TES-associated binding sites are occupied by MYC following its induction is not random; rather, in both cases, it occurs sequentially at sites that are increasingly remote from these landmarks. Conversely, MYC vacates these sites in the reverse order when its levels decline (27). This could reflect differences in the sites’ intrinsic affinities and/or in the order in which they become accessible during the course of transcriptionally permissive chromatin modification and subsequent reversion. Fourth, DNA binding around TSSs and TESs markedly stabilizes MYC, increasing its mean half-life from the standardly accepted 20-30 min to >24-48 hr (27, 72). Lastly, in addition to MYC and/or MAX, numerous other TFs reside in close proximity of TSSs and TESs and, in doing so, create unique local “neighborhoods” that are readily distinguishable from one another and from those at more remote enhancers (27).

In light of these differences, we correctly surmised that genes harboring different MYC and RNAPII footprint combinations at TSS- and TES-proximal sites would also show distinct distributions of other TF binding motifs. Indeed, this was the case for all eight alphabetical categories (Figures 2C and 3, Supplementary Files 3-10). This suggested that the effects of MYC and RNAPII on gene expression are also likely to be dictated by the chromatin landscapes generated by the integrated binding of additional TFs, their proximities to one another and their collective influence on transcription (27, 73). These might affect MYC and/or RNAPII binding and activity by virtue of non-mutually exclusive interactions involving steric hindrance, competitive binding to the same sites, or epigenetic modifications. The extent to which these interactions impact one another could also be influenced by the affinities of the TFs for their own respective DNA sites as we have documented in the case of MYC and MAX themselves (27). Variations in E box flanking sequences, i.e. the “extended motifs” that dictate the identities and binding affinities of TFs such as USF1, CLOCK and ARNT, and all of which compete with MYC-MAX and MXD-MAX heterodimers, are potential sources of yet additional transcriptional and read-through variability (13, 55, 56, 74) (Supplementary Figure 2). Differences in the abundance of these E box variants at TSSs and TESs may further explain the latter site’s lower mean affinities for MYC and MAX (27). Similarly, subtle sequence differences in SP1/3 and KLF family motifs may also determine the degree to which MYC-MAX heterodimers in association with SP1/3 are able suppress target gene expression as well (62). The presence of multiple TF binding motifs and their associated protein footprints (27), is reminiscent of promoters and enhancers, many of which can bind as many as 50 different TFs in dynamic and modular ways that confer subtle and bespoke control over gene expression (33, 75).

MYC and RNAPII cooperatively bind at TSSs and at nearby downstream RNAPII pause sites but usually dissociate rapidly as transcription resumes beyond the latter points (7, 8, 36, 39). Thus MYC’s presence at TES-proximal regions most likely represents its re-engagement with the 3’-end of the gene whereas RNAPII binding alone may be more attributable to the vestiges of processive transcription and secondary pausing (8, 37, 64, 76). This is supported by the fact that MYC and RNAPII footprints at TES-proximal sites seldom overlap precisely. As a result, the DNA motifs associated with each factor’s peak summits are generally dissimilar (alphabetical cats. d and h) (Figure 2B and D). In the case of MYC, the fact that many of these motifs are also E boxes (Figure 2 D) further supports the idea that TES-associated binding need not require or be preceded by TSS binding. Category switching provides yet additional support for this model (Figure 5A-C).

Cats. 2,6, and 13 lacked any detectable RNAPII binding at either TSSs or the usual nearby downstream pause sites but were nonetheless highly expressed and bound MYC and/or RNAPII around TESs (Figure 4A-C). Non-mutually exclusive factors that could account for this include TSS-associated binding that was functionally productive but below the threshold level of detection, rapid progression through the gene immediately upon binding and with no specific sites of pausing, or only occasional binding to accommodate genes whose transcripts are extremely stable (77, 78). Regardless of the reasons for this absence, these categories were distinct from those that did obviously bind RNAPII at TSSs (cats. 8,7,and 15) (Figure 4D) in at least three ways. First, on average, the latter were expressed at 2.1-fold higher levels than the former (Figure 4B and D). Second, they were enriched for different subsets of functionally-related genes (Figure 6A). Lastly, clear cell type-specific differences were noted in GSEA. Thus, whether the differences in RNAPII at TSSs were qualitative or quantitative in nature, their inclusion into distinct categories appears justified.

Aside from their interactions with TSSs, TESs and enhancers (27), it is possible that MYC and other TFs also regulate gene expression via other means. For example, as many as half of all TFs, including MYC, bind to and affect the trafficking, stability, and translational efficiency of mRNAs (79–81). MYC’s binding sites in these different transcripts tend to be short 5’-end regions of high GC, GA, and GU content (79). Binding recruits the mRNAs to their respective gene promoters, increases MYC’s association with non-canonical E box sites located there and creates positive transcriptional feedback loops (79). Many of the top motifs identified in the vicinity of TES-associated MYC footprints contain clusters of GA-rich motifs for TFs such as IZKF3, ELK4A, and ETV6 with which MYC could potentially interact in mRNAs (Figures 2D and 3). Depending upon whether these sites reside upstream or downstream of TESs and are incorporated into standard or DoG+ transcripts (Figure 1), they might support functions that are distinct from but cooperate with those that are dependent upon DNA binding. The concentration of MYC and other TFs at double-stranded DNA sites during active transcription might also facilitate their transfer to RNA during transcription-coupled DNA strand separation. The exchange of MYC between DNA and mRNA could also represent one way by which category switching is linked to and coordinated with gene expression. The direct transfer of MYC from the 5’- or 3’-ends of genes to transcripts might be facilitated as a direct consequence of its mediating the previously observed direct contact between TSSs and TESs (27).

The number of genes with TES-associated MYC and/or RNAPII binding that we have reported here and elsewhere is largely based on steady state levels in cell lines and tissues (27). As such, they undoubtedly underestimate the extent, plasticity and importance of category switching and the fraction of genes capable of such binding. For example, many cat. 9 genes are observed in serum-stimulated NIH3T9 fibroblasts that would otherwise not be recognized as undergoing category switching had only the initial and final time points been examined (Figure 5C upper brackets). In fact, an analysis of several intermediate time points revealed numerous instances of transient TES-associated MYC and RNAPII binding. Less dramatic but still prominent changes of a similar nature were seen in a single intermediate time point in LPS-treated B cells (Figure 5B). Others could almost certainly be identified had additional time points been examined. Similar category switching also likely occurs in murine livers during the course of HCC induction or regression (Figure 5A), in murine fibroblasts in response to doxycycline-induced MYC induction and in other cancer cells following MYC’s pharmacologic inhibition (27, 65). Category switching involving transient changes at TESs is thus likely to be quite common among genes whose transcription must be tightly, rapidly, and/or coordinately regulated, particularly in response to proliferation-related demands.

Just as TFs that localize to TSSs and enhancers influence the choice of alternative TESs so too did the binding of MYC and RNAPII around TESs regulate DoG transcription (Figure 7A) (82–84). However, there was no obvious consistency in the contributions made by these factors. Moreover, the small differences which they did exert on fractional DoG transcription appeared to be highly cell type-specific and were much less than their effects on overall transcription (Figure 4A). Together, these findings suggest that factors other the MYC and RNAPII are involved in determining whether and the degree to which genes are engaged in DoG transcription. Although normal and stress-related DoG transcription can clearly be regulated by factors such as MYC binding as TESs, an equally or more important function of this binding (along with that of RNAPII) may be to mark and coordinate the expression of genes with common functions (Figure 7C-F).

Given our earlier findings that MYC and/or MAX binding at TES-proximal sites is associated with an “open” and transcriptionally permissive local chromatin landscape, it seems reasonable to assume that this also favors DoG transcription, particularly when RNAPII is already present at these sites as it is in cats. 4,6,7 and 11-15) (Figure 2B) (27). MYC and MAX-bound TESs, much more so than sites lacking this binding, can more readily engage in direct physical contact with TSSs via “looping”, which is abolished when the E boxes underlying the TES footprints are eliminated (27). We have proposed a model by which TSS-TES contacts, possibly aided or stabilized by additional interactions with enhancers, increase the efficiency and rate of transcriptional re-initiation (27). Simultaneously, this configuration offers an option for transcription to continue beyond its standard termination site, thereby increasing the efficiency of DoG transcription while controlling total transcript levels as well. Our current findings agree with those of others showing that stresses such as hypoxia, hyperthermia and oxidative and osmotic shock cause proliferative arrest (and its attendant down-regulation of MYC); reduced RNAPII both globally and at TESs; and altered levels of DoG transcription (29–32, 85–87). It will be important in future work to explore the means by which these factors select for functionally enriched gene sets, regulate the quality and quantity of DoG transcription and identify and mark functionally related genes. Additional questions that may provide insights into more nuanced aspects of how MYC and RNAPII binding near TESs impacts transcription include the number of such sites; their distance from one another; binding by other TFs, including SP1/3, ZBTB17 and MXD members; and the degree to which these factor facilitate or inhibit direct physical interactions with TSSs or enhancers(73).

## Materials and Methods

### Accession and annotation of ChIPseq data

binding sites of MYC to those of RNAPII using the venn function from the purrr package MYC and RNAPII ChIPseq and RNAseq results for the indicated human and murine cell lines and tissues were obtained from the ENCODE database and previously published findings (https://www.encodeproject.org/) (Supplementary Table 10) (27, 66–68). Data were imported into RStudio Version 4.4.0 (http://www.posit.co/) on the University of Pittsburgh’s Computing Cluster. ChIPseq binding peaks obtained in the IDR threshold format were transformed into the narrowPeak format and finally to a Granges object using the ChIPpeakAnno package from Bioconductor (88). Granges peaks were annotated using the annotatePeakinBatch function based on their genomic coordinates, with the GRCh38 and mm10 builds being used for human and mouse data, respectively. Peaks were considered to be TSS-associated or TES-associated if their summit mapped to within +/- 2.5 kb of the TSS or TES, as previously described (27). Any genes with TSSs and TESs within 2.5 kb of one another were excluded due to site assignment ambiguity as were genes adjacent to one another in “head-to-tail” orientation (27). After annotation, genes were grouped in R using the dplyr, tidyr, stringr, and tidyverse packages based on the binding patterns of MYC and RNAPII. Locations of binding sites were plotted using the ggplot2 package. As per convention, positive values indicate binding downstream of the gene element and negative values to indicate binding upstream. To classify peaks into alphabetical categories a-h (Figure 2B), we compared the in RStudio. We considered MYC and RNAPII to be co-bound at a particular site when their signals overlapped by ≥100 base pairs (bps) (4). Each set of peaks was anti-joined to the others in Rstudio (dplyr package) to ensure that there were no accidental duplicates or shared peaks across categories.

To identify consensus TF binding sites associated with MYC and RNAPII ChIP peaks, we first performed Analysis of Motif Enrichment (AME) (MemeSuite Tools). For this analysis, we considered MYC and RNAPll peaks in categories a-h. We isolated regions +/- 50 bps on either side of the peak summits. These conservatively chosen sequences (51) were reformatted into a .bed file containing the raw genomic coordinates of the boundaries. Using bedtools v2.31.1 run from the command line, .bed files were converted into fasta files by comparison with the hg38 or mm10 genome for human and mouse cell lines. The hg38 genome (October 2022 version) was downloaded from the UCSC website (https://hgdownload.gi.ucsc.edu/goldenPath/hg38/bigZips/latest/), The mm10 genome (April 2021 version) was downloaded similarly from the UCSC website (https://hgdownload.gi.ucsc.edu/goldenPath/mm10/bigZips/latest/). We imported JASPAR (2024) CORE non-redundant position frequency matrices for vertebrates in MEME file format (jaspar.elixir.no/downloads/) and generated a master list of 879 consensus TF binding motifs. AME v5.8.8 was then used to compare output fasta files with the motif list from the JASPAR database. As a control, random shuffling of the primary input sequences was used. Output files were imported into Rstudio. The most significant 25% of results for each output file was retained while the remaining data were filtered. -log(P values) for the remaining results were calculated and plotted using the ComplexHeatmap package in Rstudio to create Fig. 2C. Because row clustering was used for these graphs, the order of motifs shown on heat maps could not be directly compared among cell lines. From these results, we selected the top TF binding motifs associated with MYC and RNAPII summits based on P values, E values (which consider true positive rates, true/false positive rates, and motif length/complexity) (Figure 2C and Supplementary Files 3-10). Some motifs, such as those for SP1 and ZBTB17, were not among the top 20-25 but were included because of the known association of their cognate TFs with MYC-MAX heterodimers (14–16); in such cases, the ranking of these sites is indicated. Following the above rank ordering, we next used FIMO (Find Individual Motif Occurrences) analysis with the above selected motifs. For this, the original JASPAR position frequency matrix files were filtered to contain only the selected motifs. The same fasta sequences used in the AME analysis were then imported into FIMO v5.5.8 and a threshold P value of 10^-4^ was established for this analysis. FIMO was run and output files were imported into RStudio for analysis. The -log_10_(P value) of every result was calculated and plotted using the ComplexHeatmap package, similar to the AME analysis. Because these results were not computationally clustered, comparisons between cell lines were made possible. FIMO uses stricter criteria to identify motifs than does AME. Thus, at times, a sequence identified as being one of the top 20 most enriched motifs by AME analysis could not be confirmed by FIMO, in which case it was replaced with the motif next in line (Figure 2D).

### Analysis of RNAseq data

RNAseq data from ENCODE or elsewhere (66–68) were imported into RStudio Version 4.4.0 directly (Supplementary Table 10), with the raw data acting as our processed data. Non-ENCODE GEO datasets included GSE76062 for livers and HCC, GSE126338 for B cells and GSE98420 for 3T9 fibroblasts. Genes having TPM values <0.1 were removed from all datasets. Previously isolated gene groups, determined by their binding s, were extracted from the data. Mean TPM values were calculated using base R, and error bars were calculated as the standard deviation of the TPM values divided by the square root of the sample size. Plots were then constructed using the ggplot2 package. P values were computed by running a two-sided, Student’s t-test between each pairwise group in the dataset.

Following peak analysis, genes were assigned to the alphanumerical categories shown in Figure 2B using the venn (purr package) intersect (BioConductor package) functions. For this, the mean and standard deviation of TPM values were calculated with data obtained from the ENCODE data base. Raw TPM values were used as post-processed data without any further corrections. For each possible comparison, a P value was calculated using an unpaired t-test with a significance cutoff of P=0.05. P value matrices were constructed and the –log_10_ values were plotted as heatmaps in R (University of Pittsburgh Computing Resources R Version 4.4.0). All non-significant comparisons were depicted in gray.

### Functional characterization of transcript categories 1-16

Transcripts from each category were obtained as previously described (27). Transcript versions were removed and subsequently mapped to ENTREZ Gene identifiers using Ensembl BioMart R package. The genes were then screened against the enrichGo database using the compareCluster function in the clusterProfiler package (version 4.4.0) (89). Human transcripts were queried against the hsapiens_gene_ensembl dataset, and mouse transcripts against mmusculus_gene_ensembl. Only unique, non-missing Entrez Gene identifiers were retained for enrichment analysis. When using the compareCluster function, our analysis included all three subsets of the GO database, namely biological process, cellular component, and molecular function. Resulting P-values were corrected using the Benjamini–Hochberg procedure to Q-values. The resulting Q-values were filtered for Q <0.05 and a GO term-by-cluster matrix was constructed. For heatmap visualization purposes, all non-significant entries were set to 0 and all entries that fit -log_10_ (Q-val) > 5 were capped at 5 for visualization purposes. Using the pheatmap package in R, a heatmap of the significance of GO terms for each cell line category combination was generated. Rows (GO terms) were hierarchally clustered.

For the category specific analysis, we used the data from above (the composite heatmap). For each GO term, the ’mean’ enrichment score, -log_10_ (Q-val), across all seven cell lines was calculated. The 20 most enriched GO terms were isolated for each category and re-plotted as heatmaps with only those selected terms. Hierarchal clustering was again used for rows. Cell lines which showed no significant enrichment in any of the 20 terms were removed from the heatmap for visualization purposes.

### Quantification of DoG transcription and functional comparisons

DoG transcription was initially assessed as previously described (27). Only protein-coding genes were used for the analysis since lncRNAs tend to have poorly defines sites of transcriptional termination(90) Briefly, BAM files were generated from raw ENCODE, rRNA-depleted strand specific RNA datasets (Supplementary Table 10). We tested two different gene boundaries: those corresponding to stranded and unstranded RNAs. Readthrough was considered as the read count greater than 500 bp downstream from the TES and total expression was considered as the read count 500 bp upstream of the TES. For each cell line, we averaged the readthrough value from two replicates (separate ENCODE experiments). We only considered protein coding genes for this analysis because of the poor definition of a TES region for other types of genes such as lncRNAs. We calculated readthrough as a ratio between the read count and the total expression. Transcripts originating from cats. 1-15 in each cell line were imported and transformed back to their original genes. We excluded cat. 16 genes because of the extremely low expression of their transcripts. This low expression made it impossible to set a threshold for meaningful read-through against data set noise. The 15 transformed gene groups were concatenated down to four for each cell line: genes with no MYC or RNAPll, those with only MYC, those with only RNAPll, and those with both at the TES. For each cell line, we plotted the read-through value distribution as a violin plot which contains a box plot within it using the ggplot2 package. We compared groups within each cell line using the Mann Whitney U Test. We chose this test after careful consideration of multiple mean and median-based tests. However, because of the vastly different group sizes and skewed data, we found mean-based tests such as Welch’s t-test to be overpowered. Contrarily, the Mann Whitney test was underpowered as it did not always detect ‘obvious’ differences. As compensation, we decided to choose the underpowered Mann Whitney U Test but set the significance threshold at p=0.10.

To compare the gene set enrichment between DoG transcription + and - genes, the above data were again concatenated, this time across cell lines to create a ‘human’ group and a ‘mouse’ group. The gene labels from each TF-pattern species group were screened against the enrichGo database using the same compareCluster function (Benjamin-Hocking p-value adjustment, p-val cutoff = 0.05, all three ontologies). Results were modified using a -log_10_(q-value) transformation. Initially, hierarchal row clustering was used to plot a heat map and reveal overarching differences between DoG transcription + and - groups. The twenty *most* enriched functional labels from each of the positive readthrough groups were then selected based on the modified E-values and plotted on separate heat maps without row clustering. All heatmaps pertaining to functional categorization of DoG transcription + gene groups were created using the ComplexHeatmap package.

## Acknowledgements

This research was supported in part by the University of Pittsburgh Center for Research Computing. Specifically, this work used the HTC cluster, which is supported by NIH award number S10OD028483. The work was also supported by Award No. 22N42 from The Rally Foundation for Childhood Cancer Research, by Innovation Grant No. 1419786 from The Alex’s Lemonade Stand Foundation for Childhood Cancer and by The UPMC Children’s Hospital Foundation (all to EVP).

## Author Contributions

C.M.H. and H.W.: performed research, analyzed data; C.M.H, H.W and E.V.P.: designed research and wrote the paper.

## Supporting information

### Supplementary Figure Legends

**Supplementary Figure 1.**
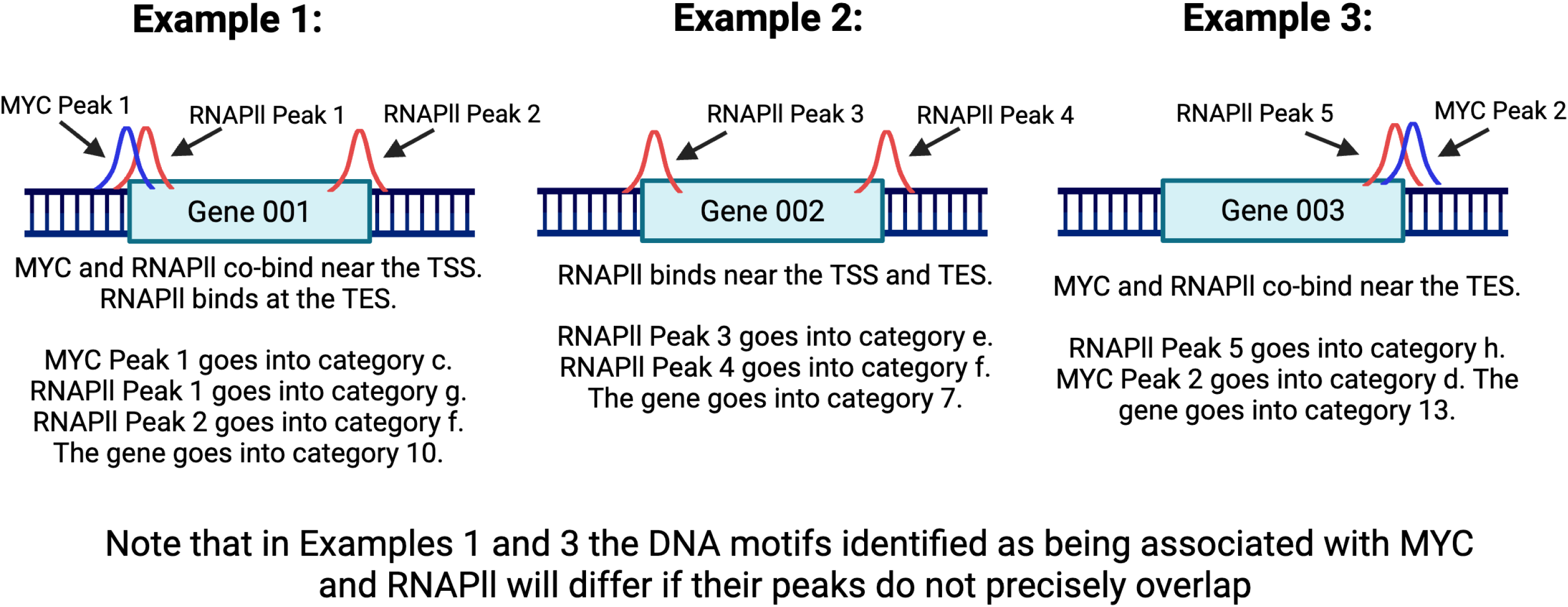
Method for classification of genes into individual alphanumeric categories based on sites and overlap of ChIPseq peaks. This scheme was used to place genes into the categories depicted in Figure 2B. Three examples (1–3) are provided here to illustrate how different categories were determined based upon the location and overlap of MYC and RNAPII ChIPseq peaks.

**Supplementary Figure 2.**
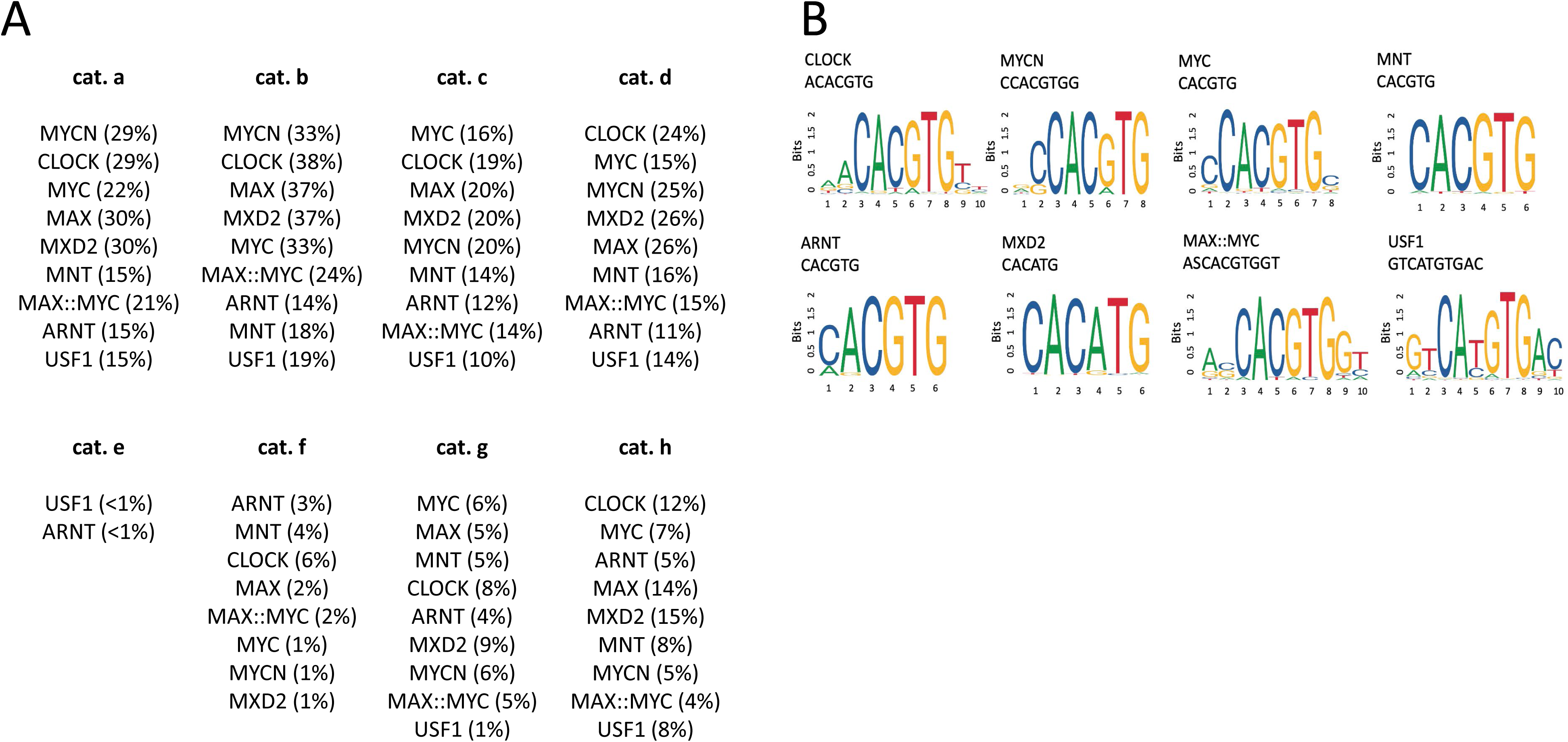
Specific motifs classified as generic E boxes in Figure 2C and. **D.** (A). Frequencies with which generic E box motifs were sub-classified into specific subsets at MYC and RNAPII-associated TSSs and TESs (C) Homer plots of the specific consensus sites used to classify various E box subsets

### Supplementary Table Legends

**Supplementary Table 1.**
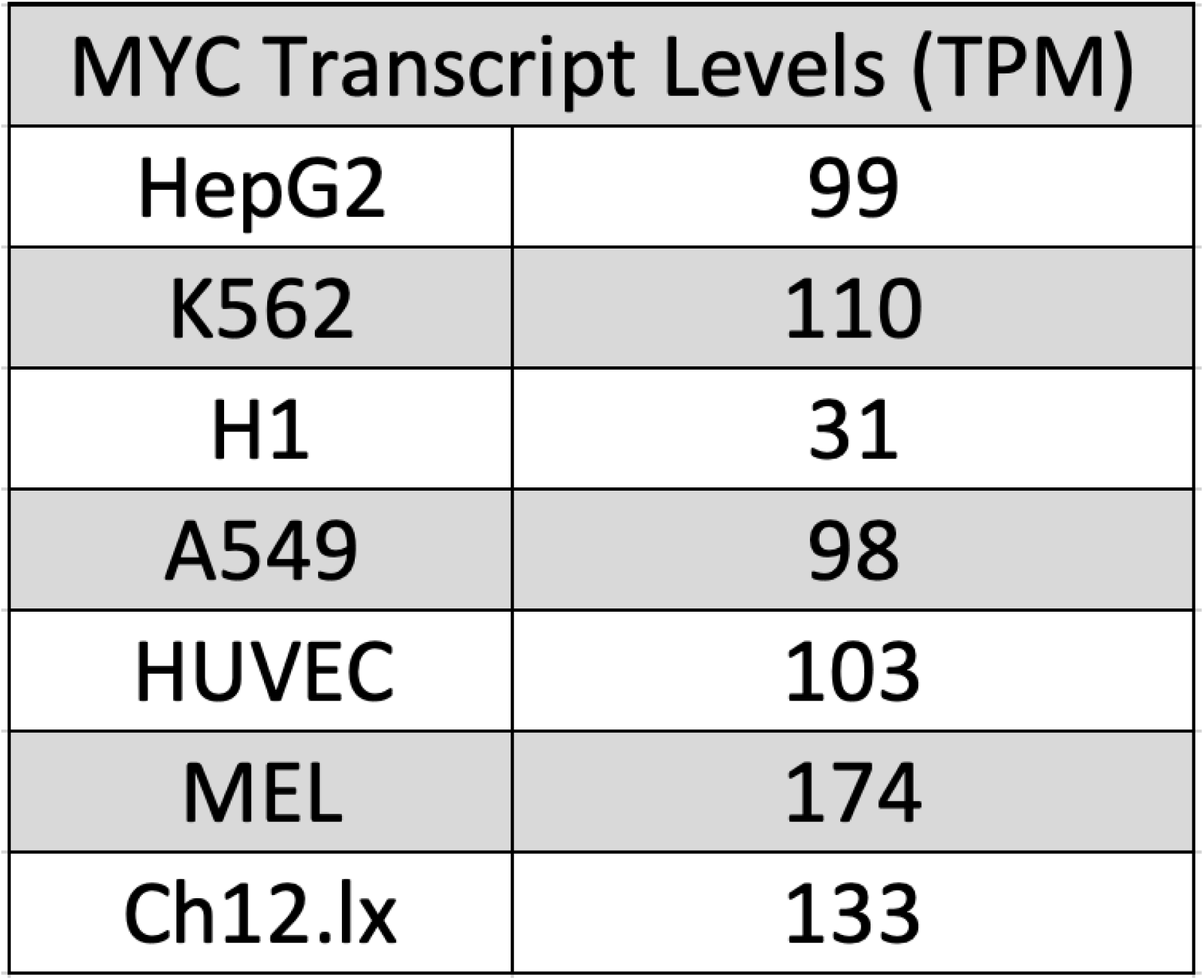
Levels of MYC transcript expression (TPM) in the indicated cell lines.

**Supplementary Table 2.** Proximity of Myc and RNAPIIll footprint peaks to TSSs and TESs.

|  | Cell Line | Distance of footprint peak summit from TSS (bps) (%) |  |  | Distance of footprint peak summit from TES (bps) (%) |  |  |
| --- | --- | --- | --- | --- | --- | --- | --- |
|  |  | 0-250 bp | 251-500 bp | > 500 bp | 0-500 bp | 501-1000 bp | > 1000 bp |
| MYC | HepG2 | 43.8 | 18.3 | 37.9 | 18.7 | 19.7 | 61.6 |
|  | K562 | 45.6 | 15.4 | 39.0 | 20.3 | 20.6 | 59.2 |
|  | H1 | 49.8 | 18.2 | 32.0 | 21.7 | 18.9 | 59.4 |
|  | A549 | 48.4 | 18.6 | 33.0 | 19.6 | 19.6 | 60.8 |
|  | HUVEC | 49.8 | 16.8 | 33.4 | 20.2 | 20.2 | 59.6 |
|  | MEL | 33.8 | 16.7 | 49.5 | 18.1 | 19.4 | 62.6 |
|  | CH12.lx | 37.0 | 16.2 | 46.8 | 17.2 | 19.1 | 63.7 |
|  | Average +/- SE | 44.0 +/- 2.4 | 17.2 +/- 0.5 | 38.8 +/- 2.6 | 19.4 +/- 0.6 | 19.6 +/- 0.3 | 61.0 +/- 0.6 |
| RNAPII | HepG2 | 45.1 | 16.0 | 38.8 | 19.5 | 22.1 | 58.4 |
|  | K562 | 43.5 | 21.6 | 34.9 | 19.9 | 24.0 | 56.0 |
|  | H1 | 46.6 | 18.0 | 35.4 | 17.6 | 23.1 | 59.3 |
|  | A549 | 55.4 | 17.1 | 27.5 | 19.7 | 22.6 | 57.7 |
|  | HUVEC | 50.7 | 15.6 | 33.7 | 17.6 | 21.6 | 60.8 |
|  | MEL | 57.1 | 13.8 | 29.0 | 19.0 | 24.5 | 56.5 |
|  | CH12.lx | 59.0 | 13.2 | 27.8 | 17.8 | 23.2 | 59.0 |
|  | Average +/- SE | 51.1 +/- 2.3 | 16.5 +/- 1.0 | 32.5 +/- 1.6 | 18.7 +/- 0.4 | 23.0 +/- 0.4 | 58.2 +/- 0.7 |

**Supplementary Table 3.**
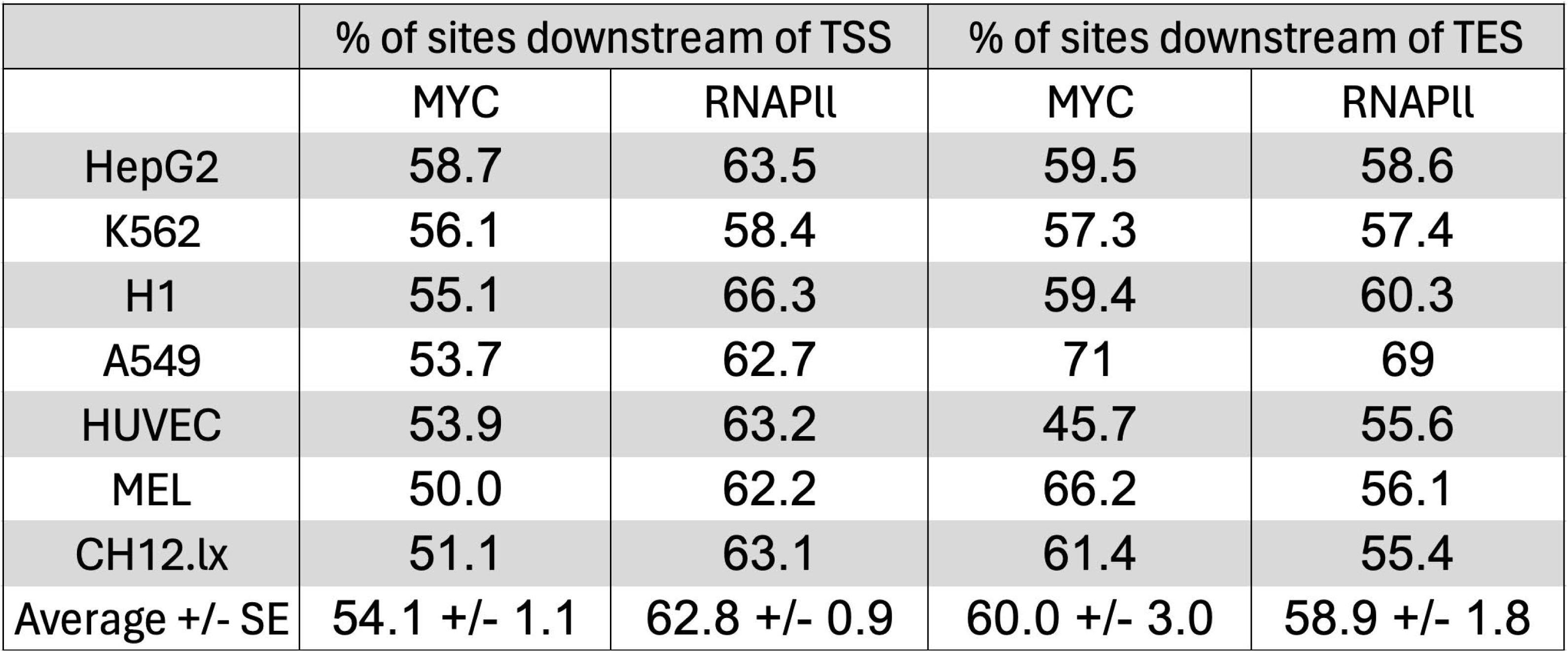
Binding sites distributions of MYC and RNAPII around TSSs and TESs in genes that bind both factors.

**Supplementary Table 4.** Average distance (in bps) of MYC and RNAPII binding peak summits from TSSs.

|  | MYC Peak Summit Distance |  |  | RNAPII Peak Summit Distance |  |
| --- | --- | --- | --- | --- | --- |
|  | Peak 1 | Peak 2 |  | Peak 1 | Peak 2 |
| MYC and RNAPII Binding |  |  | MYC and RNAPII Binding |  |  |
| HepG2 | 10 | NA | HepG2 | -220 | 70 |
| K562 | 0 | NA | K562 | -250 | 90 |
| H1 | -20 | NA | H1 | 60 | NA |
| A549 | -30 | NA | A549 | -250 | 70 |
| HUVEC | -20 | NA | HUVEC | 70 | NA |
| MEL | -50 | NA | MEL | -160 | 70 |
| CH12.lx | 40 | NA | CH12.lx | -180 | 60 |
| Average +/- SE | -10.0 +/- 11.1 | NA | Average +/- SE | -132.9 +/- 52.6 | 72 +/- 4.9 |
| MYC Binding Only |  |  | RNAPII Binding Only |  |  |
| HepG2 | 0 | NA | HepG2 | 70 | NA |
| K562 | -40 | NA | K562 | -220 | 110 |
| H1 | -40 | NA | H1 | -150 | 60 |
| A549 | -70 | NA | A549 | -150 | 60 |
| HUVEC | -60 | NA | HUVEC | -170 | 60 |
| MEL | -110 | NA | MEL | -140 | 60 |
| CH12.lx | -210 | NA | CH12.lx | -140 | 50 |
| Average +/- SE | -75.7 +/- 25.7 | NA | Average +/- SE | -128.6 +/- 34.7 | 66.7 +/- 8.8 |

**Supplementary Table 5.** Average distance (in bps) of MYC and RNAPII binding peak summits from single or multiple TESs.

|  | MYC Peak Binding Sites |  |  |  | RNAPII Peak Binding Sites |  |  |
| --- | --- | --- | --- | --- | --- | --- | --- |
|  | Peak 1 | Peak 2 | Peak 3 |  | Peak 1 | Peak 2 | Peak 3 |
| MYC and RNAPII Binding |  |  |  | MYC and RNAPII Binding |  |  |  |
| HepG2 | -700 | 500 | 1250 | HepG2 | -2200 | 500 | 1750 |
| K562 | -1100 | 680 | NA | K562 | 540 | 2170 | NA |
| H1 | -1010 | 170 | 1620 | H1 | -960 | 990 | NA |
| A549 | -1150 | 1100 | NA | A549 | 1200 | NA | NA |
| HUVEC | -500 | NA | NA | HUVEC | -600 | 1650 | NA |
| MEL | -1500 | 900 | NA | MEL | -1600 | -500 | 900 |
| CH12.lx | -2100 | 900 | NA | CH12.lx | -2300 | -500 | 850 |
| Average +/- SE | -1151.4 +/- 500.0 | 708.3 +/- 126.6 | 1435 +/- 101.1 | Average +/- SE | -845.7 +/- 504.7 | 718.3 +/- 416.3 | 1166.7 +/- 191.2 |
| MYC Binding Only |  |  |  | RNAPII Binding Only |  |  |  |
| HepG2 | -2100 | -580 | 575 | HepG2 | -660 | 650 | NA |
| K562 | -1870 | 480 | 2110 | K562 | 760 | NA | NA |
| H1 | -1800 | 525 | NA | H1 | -670 | 660 | NA |
| A549 | -1890 | 570 | 1800 | A549 | 660 | NA | NA |
| HUVEC | 150 | 1910 | NA | HUVEC | 890 | NA | NA |
| MEL | -1900 | 600 | 1880 | MEL | -540 | 700 | NA |
| CH12.lx | -1830 | 570 | 1830 | CH12.lx | 710 | NA | NA |
| Average +/- SE | -1605.7 +/- 295.1 | 582.1 +/- 273.1 | 1655.0 +/- 232.1 | Average +/- SE | 164.3 +/- 280.1 | 670.0 +/- 10.0 | NA |

**Supplementary Table 6.**
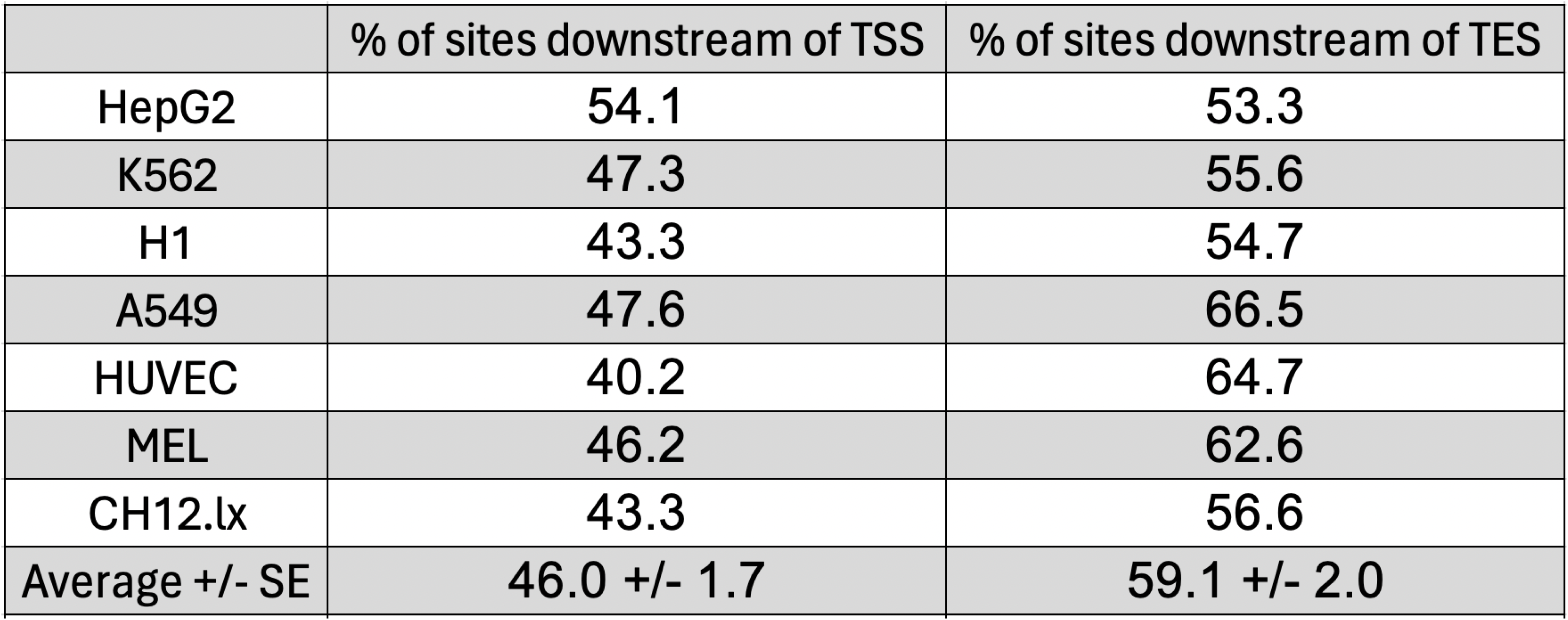
Binding site distributions of MYC around TSSs and TESs in genes that do not bind RNAPII.

|  | % of sites downstream of TSS | % of sites downstream of TES |
| --- | --- | --- |
| HepG2 | 54.1 | 53.3 |
| K562 | 47.3 | 55.6 |
| H1 | 43.3 | 54.7 |
| A549 | 47.6 | 66.5 |
| HUVEC | 40.2 | 64.7 |
| MEL | 46.2 | 62.6 |
| CH12.lx | 43.3 | 56.6 |
| Average +/- SE | 46.0 +/- 1.7 | 59.1 +/- 2.0 |

**Supplementary Table 7.**
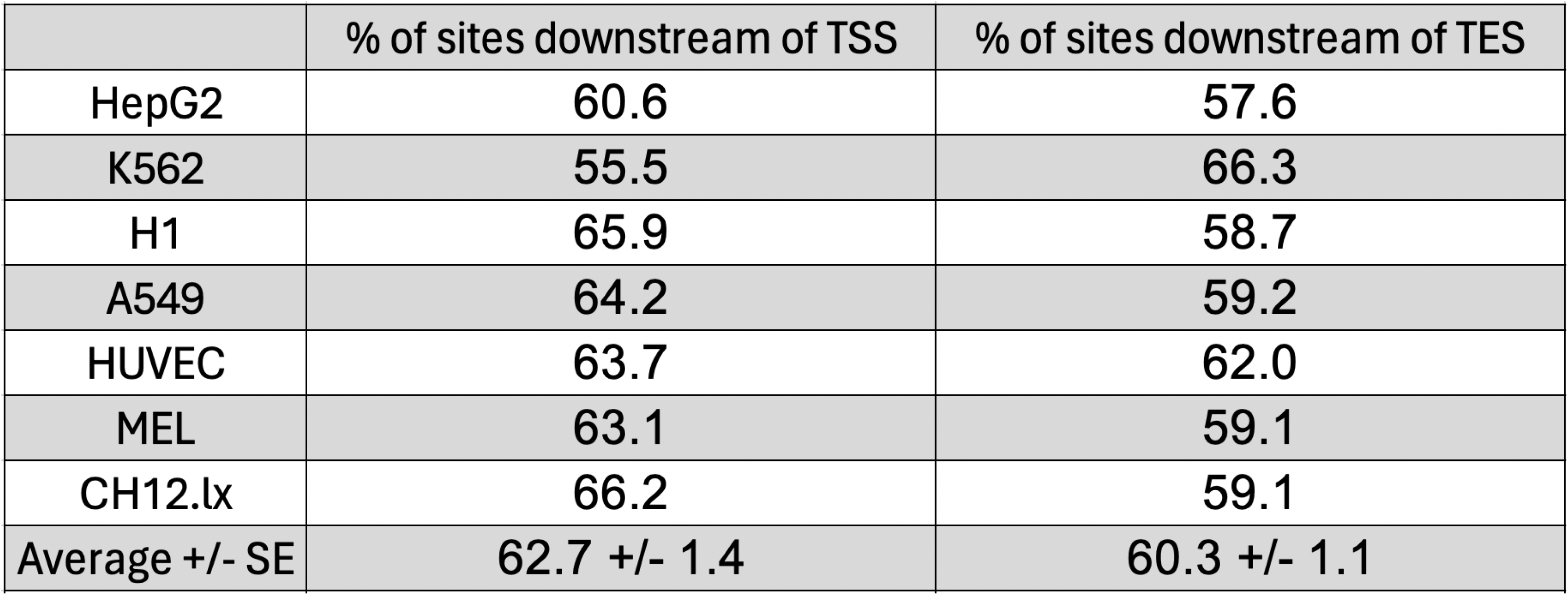
Binding sites distributions of RNAPII around TSSs and TESs in genes that do not bind MYC.

|  | % of sites downstream of TSS | % of sites downstream of TES |
| --- | --- | --- |
| HepG2 | 60.6 | 57.6 |
| K562 | 55.5 | 66.3 |
| H1 | 65.9 | 58.7 |
| A549 | 64.2 | 59.2 |
| HUVEC | 63.7 | 62.0 |
| MEL | 63.1 | 59.1 |
| CH12.lx | 66.2 | 59.1 |
| Average +/- SE | 62.7 +/- 1.4 | 60.3 +/- 1.1 |

**Supplementary Table 8.** Mean expression levels of numerical gene categories in individual cell lines.

| Cell Line | Cat 1 | Cat 2 | Cat 3 | Cat 4 | Cat 5 | Cat 6 | Cat 7 | Cat 8 | Cat 9 | Cat 10 | Cat 11 | Cat 12 | Cat 13 | Cat 14 | Cat 15 | Cat 16 |
| --- | --- | --- | --- | --- | --- | --- | --- | --- | --- | --- | --- | --- | --- | --- | --- | --- |
| HepG2 | 2.84 | 2.28 | 1.92 | 12.68 | 8.45 | 20.53 | 25.46 | 3.47 | 9.83 | 9.17 | 32.40 | 18.69 | 3.99 | 2.60 | 8.88 | 0.67 |
| K562 | 4.32 | 2.44 | 3.74 | 17.09 | 6.36 | 14.77 | 9.57 | 8.17 | 11.38 | 13.65 | 35.76 | 15.79 | 4.21 | 2.30 | 13.27 | 0.70 |
| H1 | 6.95 | 2.77 | NA | 62.20 | 11.28 | 22.67 | 38.39 | 6.94 | 15.69 | 21.21 | 61.17 | 16.82 | 6.59 | NA | 21.91 | 1.09 |
| A549 | 9.89 | 7.44 | 0.52 | 7.40 | 11.66 | 6.29 | 14.48 | 11.39 | 21.21 | 37.61 | 76.52 | 24.07 | 4.04 | 9.89 | NA | 1.08 |
| HUVEC | 8.49 | 5.85 | 0.93 | 42.72 | 11.76 | 17.28 | 35.06 | 20.48 | 16.06 | 14.54 | 48.47 | 50.47 | 7.64 | 14.05 | 17.71 | 0.94 |
| MEL | 2.19 | 1.72 | 2.47 | 18.20 | 3.35 | 9.78 | 9.04 | 2.67 | 5.00 | 4.53 | 17.09 | 6.82 | 2.57 | 3.56 | 5.83 | 5.97 |
| CH12.lx | 1.82 | 1.23 | 1.43 | 19.94 | 3.17 | 7.31 | 8.83 | 2.55 | 4.99 | 4.12 | 17.57 | 5.88 | 3.01 | 6.93 | 4.06 | 0.23 |
| Average | 5.22 | 3.39 | 1.83 | 25.75 | 8.00 | 14.09 | 20.12 | 7.95 | 12.02 | 14.97 | 41.28 | 19.79 | 4.58 | 6.55 | 11.94 | 1.53 |

**Supplementary Table 9.** Number of surveyed genes used to determine mean expression levels in numerical categories in individual cell lines.

| Cell Line | Cat 1 | Cat 2 | Cat 3 | Cat 4 | Cat 5 | Cat 6 | Cat 7 | Cat 8 | Cat 9 | Cat 10 | Cat 11 | Cat 12 | Cat 13 | Cat 14 | Cat 15 | Cat 16 | Total |
| --- | --- | --- | --- | --- | --- | --- | --- | --- | --- | --- | --- | --- | --- | --- | --- | --- | --- |
| HepG2 | 2395 | 1198 | 77 | 33 | 1569 | 847 | 57 | 31 | 4425 | 86 | 130 | 66 | 411 | 22 | 16 | 19302 | 30665 |
| K562 | 2817 | 1666 | 71 | 28 | 1545 | 591 | 31 | 35 | 4003 | 98 | 82 | 32 | 380 | 31 | 9 | 18372 | 29791 |
| H1 | 652 | 282 | 2 | 27 | 4959 | 1688 | 244 | 12 | 1350 | 3 | 105 | 5 | 74 | 1 | 6 | 18230 | 27640 |
| A549 | 581 | 299 | 5 | 4 | 4317 | 337 | 25 | 21 | 2221 | 7 | 10 | 2 | 57 | 2 | 2 | 14830 | 22720 |
| HUVEC | 442 | 186 | 3 | 16 | 4779 | 1452 | 192 | 14 | 1148 | 3 | 56 | 3 | 118 | 1 | 13 | 19271 | 27697 |
| MEL | 1537 | 1242 | 47 | 88 | 2147 | 1121 | 121 | 74 | 3597 | 147 | 402 | 99 | 486 | 32 | 37 | 17794 | 28971 |
| CH12.lx | 1460 | 1264 | 57 | 51 | 2611 | 865 | 103 | 83 | 3852 | 151 | 272 | 109 | 489 | 24 | 45 | 18988 | 30424 |
| Total | 9884 | 6137 | 262 | 247 | 21927 | 6901 | 773 | 270 | 20596 | 495 | 1057 | 316 | 2015 | 113 | 128 | 126787 | 197908 |

**Supplementary Table 10.**
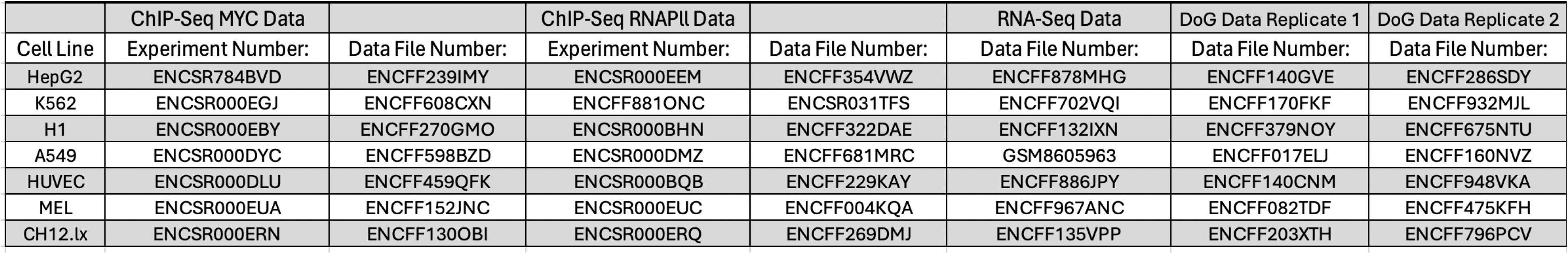
Table of raw data accession numbers from the ENCODE database for each cell line.

## Supplementary Files

**Supplementary File 1. Raw sequences compiled from the alphabetic categories a-h**

**Supplementary File 2. Raw transcript IDs compiled from the numerical categories 1-16**

**Supplementary File 3. Rank list of the most common DNA binding motifs residing within +/- 50 bps of MYC and RNAPll peak summits at TSSs and TESs across all cell lines (aggregate cell line results)**

**Supplementary File 4. Rank list of the most common DNA binding motifs residing within +/- 50 bps of MYC and RNAPll peak summits at TSSs and TESs across HepG2 cells. See Figure 3C and Supplementary File 1 for combined results**

**Supplementary File 5. Rank list of the most common DNA binding motifs residing within +/- 50 bps of MYC and RNAPll peak summits at TSSs and TESs across K562 cells. See Figure 3C and Supplementary File 1 for combined results**

**Supplementary File 6. Rank list of the most common DNA binding motifs residing within +/- 50 bps of MYC and RNAPll peak summits at TSSs and TESs across H1 cells. See Figure 3C and Supplementary File 1 for combined results**

**Supplementary File 7. Rank list of the most common DNA binding motifs residing within +/- 50 bps of MYC and RNAPll peak summits at TSSs and TESs across A549 cells. See Figure 3C and Supplementary File 1 for combined results**

**Supplementary File 8. Rank list of the most common DNA binding motifs residing within +/- 50 bps of MYC and RNAPll peak summits at TSSs and TESs across HUVEC cells. See Figure 3C and Supplementary File 1 for combined results**

**Supplementary File 9. Rank list of the most common DNA binding motifs residing within +/- 50 bps of MYC and RNAPll peak summits at TSSs and TESs across MEL cells. See Figure 3C and Supplementary File 1 for combined results**

**Supplementary File 10. Rank list of the most common DNA binding motifs residing within +/- 50 bps of MYC and RNAPll peak summits at TSSs and TESs across Ch12.lx cells. See Figure 3C and Supplementary File 1 for combined results**

**Supplementary File 11. Significantly enriched gene sets complied from Figure 6A**

**Supplementary File 12. Downstream-of-gene readthrough values compiled from Figure 7A**

**Supplementary File 13. Significantly enriched gene sets compiled from Figure 7B**

## References

1. R. Dhanasekaran et al., The MYC oncogene - the grand orchestrator of cancer growth and immune evasion. Nat Rev Clin Oncol 19, 23–36 (2022).

2. E. V. Prochownik, Regulation of Normal and Neoplastic Proliferation and Metabolism by the Extended Myc Network. Cells 11, 3974 (2022).

3. J. Sarid, T. D. Halazonetis, W. Murphy, P. Leder, Evolutionarily conserved regions of the human c-myc protein can be uncoupled from transforming activity. Proc Natl Acad Sci U S A 84, 170–173 (1987).

4. P. A. Carroll, B. W. Freie, H. Mathsyaraja, R. N. Eisenman, The MYC transcription factor network: balancing metabolism, proliferation and oncogenesis. Front Med 12, 412–425 (2018).

5. K. I. Zeller et al., Global mapping of c-Myc binding sites and target gene networks in human B cells. Proc Natl Acad Sci U S A 103, 17834–17839 (2006).

6. N. Gomez-Roman et al., Activation by c-Myc of transcription by RNA polymerases I, II and III. Biochem Soc Symp 10.1042/bss0730141, 141–154 (2006).

7. C. Lourenco et al., MYC protein interactors in gene transcription and cancer. Nat Rev Cancer 21, 579–591 (2021).

8. P. B. Rahl et al., c-Myc regulates transcriptional pause release. Cell 141, 432–445 (2010).

9. X. Qiu et al., MYC drives aggressive prostate cancer by disrupting transcriptional pause release at androgen receptor targets. Nat Commun 13, 2559 (2022).

10. E. S. Goetzman, E. V. Prochownik, The Role for Myc in Coordinating Glycolysis, Oxidative Phosphorylation, Glutaminolysis, and Fatty Acid Metabolism in Normal and Neoplastic Tissues. Front Endocrinol (Lausanne) 9, 129 (2018).

11. H. Wang et al., Disruption of Multiple Overlapping Functions Following Stepwise Inactivation of the Extended Myc Network. Cells 11, 4087 (2022).

12. E. Wolf, C. Y. Lin, M. Eilers, D. L. Levens, Taming of the beast: shaping Myc-dependent amplification. Trends Cell Biol 25, 241–248 (2015).

13. E. V. Prochownik, H. Wang, Normal and Neoplastic Growth Suppression by the Extended Myc Network. Cells 11, 747 (2022).

14. A. L. Gartel, K. Shchors, Mechanisms of c-myc-mediated transcriptional repression of growth arrest genes. Exp Cell Res 283, 17–21 (2003).

15. M. Wanzel, S. Herold, M. Eilers, Transcriptional repression by Myc. Trends Cell Biol 13, 146–150 (2003).

16. K. E. Wiese et al., The role of MIZ-1 in MYC-dependent tumorigenesis. Cold Spring Harb Perspect Med 3, a014290 (2013).

17. H. Wang et al., Premature aging and reduced cancer incidence associated with near-complete body-wide Myc inactivation. Cell Rep 42, 112830 (2023).

18. C. Y. Lin et al., Transcriptional amplification in tumor cells with elevated c-Myc. Cell 151, 56–67 (2012).

19. F. Lorenzin et al., Different promoter affinities account for specificity in MYC-dependent gene regulation. Elife 5, e15161 (2016).

20. S. Pelengaris, M. Khan, G. Evan, c-MYC: more than just a matter of life and death. Nat Rev Cancer 2, 764–776 (2002).

21. A. Sabo, B. Amati, Genome recognition by MYC. Cold Spring Harb Perspect Med 4, a014191 (2014).

22. S. T. Jakobsen et al., MYC activity at enhancers drives prognostic transcriptional programs through an epigenetic switch. Nat Genet 56, 663–674 (2024).

23. Q. Li et al., Enhancer RNAs: mechanisms in transcriptional regulation and functions in diseases. Cell Commun Signal 21, 191 (2023).

24. Y. X. See, K. Chen, M. J. Fullwood, MYC overexpression leads to increased chromatin interactions at super-enhancers and MYC binding sites. Genome Res 32, 629–642 (2022).

25. K. MacPherson-Hawthorne, R. C. Sears, Hold the MYCrophone: MYC Invades Enhancers to Control Cancer-Type Gene Programs. Cancer Res 84, 2227–2228 (2024).

26. J. Schuijers et al., Transcriptional Dysregulation of MYC Reveals Common Enhancer-Docking Mechanism. Cell Rep 23, 349–360 (2018).

27. H. Wang et al., MYC Binding Near Transcriptional End Sites Regulates Basal Gene Expression, Read-Through Transcription, and Intragenic Contacts. Adv Sci (Weinh) 12, e14601 (2025).

28. P. Caldas et al., Transcription readthrough is prevalent in healthy human tissues and associated with inherent genomic features. Commun Biol 7, 100 (2024).

29. N. A. Rosa-Mercado, J. A. Steitz, Who let the DoGs out? - biogenesis of stress-induced readthrough transcripts. Trends Biochem Sci 47, 206–217 (2022).

30. A. Vijayakumar, A. Park, J. A. Steitz, Modulation of mRNA 3’-End Processing and Transcription Termination in Virus-Infected Cells. Front Immunol 13, 828665 (2022).

31. A. Vilborg et al., Comparative analysis reveals genomic features of stress-induced transcriptional readthrough. Proc Natl Acad Sci U S A 114, E8362–E8371 (2017).

32. A. Vilborg, J. A. Steitz, Readthrough transcription: How are DoGs made and what do they do? RNA Biol 14, 632–636 (2017).

33. Y. Bilu, N. Barkai, The design of transcription-factor binding sites is affected by combinatorial regulation. Genome Biol 6, R103 (2005).

34. C. H. Chen et al., Determinants of transcription factor regulatory range. Nat Commun 11, 2472 (2020).

35. S. H. Duttke et al., Position-dependent function of human sequence-specific transcription factors. Nature 631, 891–898 (2024).

36. A. Baluapuri, E. Wolf, M. Eilers, Target gene-independent functions of MYC oncoproteins. Nat Rev Mol Cell Biol 21, 255–267 (2020).

37. V. H. Cowling, M. D. Cole, The Myc transactivation domain promotes global phosphorylation of the RNA polymerase II carboxy-terminal domain independently of direct DNA binding. Mol Cell Biol 27, 2059–2073 (2007).

38. E. J. Feris, J. W. Hinds, M. D. Cole, Formation of a structurally-stable conformation by the intrinsically disordered MYC:TRRAP complex. PLoS One 14, e0225784 (2019).

39. M. Kalkat et al., MYC Protein Interactome Profiling Reveals Functionally Distinct Regions that Cooperate to Drive Tumorigenesis. Mol Cell 72, 836–848 e837 (2018).

40. B. Turner, “ChIP with Native Chromatin: Advantages and Problems Relative to Methods Using Cross-Linked Material” in Mapping Protein/DNA Interactions by Cross-Linking. (Paris, 2001).

41. E. Wolf et al., Miz1 is required to maintain autophagic flux. Nat Commun 4, 2535 (2013).

42. M. W. Szczesniak, M. R. Kubiak, E. Wanowska, I. Makalowska, Comparative genomics in the search for conserved long noncoding RNAs. Essays Biochem 65, 741–749 (2021).

43. Y. Chen, H. Li, Y. Y. Li, Y. Li, Pan-Cancer Analysis of Head-to-Head Gene Pairs in Terms of Transcriptional Activity, Co-expression and Regulation. Front Genet 11, 560997 (2020).

44. D. R. Herr, G. L. Harris, Close head-to-head juxtaposition of genes favors their coordinate regulation in Drosophila melanogaster. FEBS Lett 572, 147–153 (2004).

45. H. Li et al., Abundant binary promoter switches in lineage-determining transcription factors indicate a digital component of cell fate determination. Cell Rep 42, 113454 (2023).

46. B. L. Barrilleaux et al., Miz-1 activates gene expression via a novel consensus DNA binding motif. PLoS One 9, e101151 (2014).

47. Y. Hasegawa, K. Struhl, Different SP1 binding dynamics at individual genomic loci in human cells. Proc Natl Acad Sci U S A 118, e2113579118 (2021).

48. K. Lysakovskaia, A. Devadas, B. Schwalb, M. Lidschreiber, P. Cramer, Promoter-proximal RNA polymerase II termination regulates transcription during human cell type transition. Nat Struct Mol Biol 32, 995–1005 (2025).

49. M. Quinodoz, C. Gobet, F. Naef, K. B. Gustafson, Characteristic bimodal profiles of RNA polymerase II at thousands of active mammalian promoters. Genome Biol 15, R85 (2014).

50. M. Commane et al., Quantitative single-cell imaging suggests increased global chromatin accessibility in tumor versus non-tumor cell lines. iScience 28, 113570 (2025).

51. X. Ma et al., A highly efficient and effective motif discovery method for ChIP-seq/ChIP-chip data using positional information. Nucleic Acids Res 40, e50 (2012).

52. P. D. Grunwald, Beyond Neyman-Pearson: E-values enable hypothesis testing with a data-driven alpha. Proc Natl Acad Sci U S A 121, e2302098121 (2024).

53. M. Allevato et al., Sequence-specific DNA binding by MYC/MAX to low-affinity non-E-box motifs. PLoS One 12, e0180147 (2017).

54. K. E. Boyd, P. J. Farnham, Myc versus USF: discrimination at the cad gene is determined by core promoter elements. Mol Cell Biol 17, 2529–2537 (1997).

55. F. Fisher et al., Transcription activation by Myc and Max: flanking sequences target activation to a subset of CACGTG motifs in vivo. EMBO J 12, 5075–5082 (1993).

56. E. V. Prochownik, M. E. VanAntwerp, Differential patterns of DNA binding by myc and max proteins. Proc Natl Acad Sci U S A 90, 960–964 (1993).

57. H. I. Swanson, J. H. Yang, Specificity of DNA binding of the c-Myc/Max and ARNT/ARNT dimers at the CACGTG recognition site. Nucleic Acids Res 27, 3205–3212 (1999).

58. W. M. Gombert, A. Krumm, Targeted deletion of multiple CTCF-binding elements in the human C-MYC gene reveals a requirement for CTCF in C-MYC expression. PLoS One 4, e6109 (2009).

59. Z. Wei et al., MYC reshapes CTCF-mediated chromatin architecture in prostate cancer. Nat Commun 14, 1787 (2023).

60. X. Sun, J. Zhang, C. Cao, CTCF and Its Partners: Shaper of 3D Genome during Development. Genes (Basel) 13, 1383 (2022).

61. R. Kalyan Sundaram, R. Radhakrishnan, B. Lim, MYC and AP-1 oncogenes synergistically bind enhancers to rewire transcription. bioRxiv 10.1101/2025.04.28.650480 (2025).

62. C. K. Kim, P. He, A. B. Bialkowska, V. W. Yang, SP and KLF Transcription Factors in Digestive Physiology and Diseases. Gastroenterology 152, 1845–1875 (2017).

63. A. Baluapuri et al., MYC Recruits SPT5 to RNA Polymerase II to Promote Processive Transcription Elongation. Mol Cell 74, 674–687 e611 (2019).

64. D. Ezer, N. R. Zabet, B. Adryan, Homotypic clusters of transcription factor binding sites: A model system for understanding the physical mechanics of gene expression. Comput Struct Biotechnol J 10, 63–69 (2014).

65. J. M. Dolezal et al., Sequential adaptive changes in a c-Myc-driven model of hepatocellular carcinoma. J Biol Chem 292, 10068–10086 (2017).

66. T. R. Kress et al., Identification of MYC-Dependent Transcriptional Programs in Oncogene-Addicted Liver Tumors. Cancer Res 76, 3463–3472 (2016).

67. A. Tesi et al., An early Myc-dependent transcriptional program orchestrates cell growth during B-cell activation. EMBO Rep 20, e47987 (2019).

68. S. de Pretis et al., Integrative analysis of RNA polymerase II and transcriptional dynamics upon MYC activation. Genome Res 27, 1658–1664 (2017).

69. G. Chovatiya et al., Cell-type-specific RNA polymerase II activity maps in intact tissues provide a gateway to mammalian gene regulatory mechanisms in vivo. Dev Cell 61, 434–451 e438 (2026).

70. J. Huang, X. Ji, Never a dull enzyme, RNA polymerase II. Transcription 14, 49–67 (2023).

71. J. van Riggelen, A. Yetil, D. W. Felsher, MYC as a regulator of ribosome biogenesis and protein synthesis. Nat Rev Cancer 10, 301–309 (2010).

72. A. S. Farrell, R. C. Sears, MYC degradation. Cold Spring Harb Perspect Med 4, a014365 (2014).

73. I. Dror, T. Golan, C. Levy, R. Rohs, Y. Mandel-Gutfreund, A widespread role of the motif environment in transcription factor binding across diverse protein families. Genome Res 25, 1268–1280 (2015).

74. R. Gordan et al., Genomic regions flanking E-box binding sites influence DNA binding specificity of bHLH transcription factors through DNA shape. Cell Rep 3, 1093–1104 (2013).

75. C. Buecker, J. Wysocka, Enhancers as information integration hubs in development: lessons from genomics. Trends Genet 28, 276–284 (2012).

76. K. Glover-Cutter, S. Kim, J. Espinosa, D. L. Bentley, RNA polymerase II pauses and associates with pre-mRNA processing factors at both ends of genes. Nat Struct Mol Biol 15, 71–78 (2008).

77. L. J. Core et al., Defining the status of RNA polymerase at promoters. Cell Rep 2, 1025–1035 (2012).

78. M. P. Swaffer et al., RNA polymerase II dynamics and mRNA stability feedback scale mRNA amounts with cell size. Cell 186, 5254–5268 e5226 (2023).

79. S. Li et al., Integrative characterization of MYC RNA-binding function. Cell Genom 5, 100878 (2025).

80. K. S. Moore, M. von Lindern, RNA Binding Proteins and Regulation of mRNA Translation in Erythropoiesis. Front Physiol 9, 910 (2018).

81. O. Oksuz et al., Transcription factors interact with RNA to regulate genes. Mol Cell 83, 2449–2463 e2413 (2023).

82. C. Alfonso-Gonzalez et al., Sites of transcription initiation drive mRNA isoform selection. Cell 186, 2438–2455 e2422 (2023).

83. B. Kwon et al., Enhancers regulate 3’ end processing activity to control expression of alternative 3’UTR isoforms. Nat Commun 13, 2709 (2022).

84. A. Reyes, W. Huber, Alternative start and termination sites of transcription drive most transcript isoform differences across human tissues. Nucleic Acids Res 46, 582–592 (2018).

85. J. F. Cardiello, J. A. Goodrich, J. F. Kugel, Heat Shock Causes a Reversible Increase in RNA Polymerase II Occupancy Downstream of mRNA Genes, Consistent with a Global Loss in Transcriptional Termination. Mol Cell Biol 38, e00181–00118 (2018).

86. S. Fulda, A. M. Gorman, O. Hori, A. Samali, Cellular stress responses: cell survival and cell death. Int J Cell Biol 2010, 214074 (2010).

87. L. Mansouri, Y. Xie, D. A. Rappolee, Adaptive and Pathogenic Responses to Stress by Stem Cells during Development. Cells 1, 1197–1224 (2012).

88. R. C. Gentleman et al., Bioconductor: open software development for computational biology and bioinformatics. Genome Biol 5, R80 (2004).

89. G. Yu, L. G. Wang, Y. Han, Q. Y. He, clusterProfiler: an R package for comparing biological themes among gene clusters. OMICS 16, 284–287 (2012).

90. J. S. Mattick et al., Long non-coding RNAs: definitions, functions, challenges and recommendations. Nat Rev Mol Cell Biol 24, 430–447 (2023).

91. M. Ronzio, F. Zambelli, D. Dolfini, R. Mantovani, G. Pavesi, Integrating Peak Colocalization and Motif Enrichment Analysis for the Discovery of Genome-Wide Regulatory Modules and Transcription Factor Recruitment Rules. Front Genet 11, 72 (2020).

92. K. Kimura, T. L. B. Jackson, R. C. C. Huang, Interaction and Collaboration of SP1, HIF-1, and MYC in Regulating the Expression of Cancer-Related Genes to Further Enhance Anticancer Drug Development. Curr Issues Mol Biol 45, 9262–9283 (2023).

